# When Cells Rebel: a comparative genomics investigation into marsupial cancer susceptibility

**DOI:** 10.1101/2025.08.12.669816

**Authors:** Cleopatra Petrohilos, Emma Peel, Luke W. Silver, Patrick G. S. Grady, Rachel J. O’Neill, Carolyn J. Hogg, Katherine Belov

## Abstract

Cancer is ubiquitous in multicellular life, yet susceptibility varies significantly between species. Previous studies have shown a genetic basis for cancer resistance in many species, but few studies have investigated the inverse: why some species are particularly susceptible to cancer. The Dasyuridae are a family of carnivorous marsupials that are frequently reported as having high rates of cancer prevalence. We hypothesised that this high susceptibility also has a genetic basis. To investigate this, we generated reference genomes for the kowari (*Dasyuroides byrnei*), a dasyurid species with one of the highest rates of reported cancer prevalence among mammals, and a non-dasyurid marsupial, the eastern barred bandicoot (*Perameles gunnii*). We used these to perform a comparative genomics analysis alongside nine previously assembled reference genomes: four dasyurid species and five non-dasyurid marsupial species. Genomes were annotated using FGENESH++ and assigned to orthogroups for input to CAFE (Computational Analysis of gene Family Evolution) analysis to identify gene families that had undergone significant expansions or contractions in each lineage. In the dasyurids, we identified large expansions in Ras genes, a family of oncogenes. Interestingly, a similar expansion of Ras genes was also identified in the bandicoot and bilby. These genes were primarily expressed in tissues such as testes, ovaries and yolk sac, so we hypothesise they serve a reproductive role. Future work is required to identify the potential roles of oncogene expansions in cancer susceptibility in these marsupial species.

## Introduction

Cancer encompasses a multitude of diseases that are all characterised by uncontrolled cell growth and metastasis (Hanahan, 2022). They arise when mutations accumulate in the three major groups of driver genes: tumour suppressors, proto-oncogenes, and DNA damage response (Arnal et al., 2016). Mutations in tumour suppressors result in a loss of function (Levitt & Hickson, 2002) whilst mutated DNA repair genes cause genome instability, accelerating the rate at which deleterious mutations accumulate (Arnal et al., 2016; Jeggo et al., 2016). Mutated proto-oncogenes cause a gain of function (Chan & Feng, 2007) and result in uncontrolled cell proliferation (Arnal et al., 2016).

Cancer is ubiquitous in multicellular life (Aktipis et al., 2015). Phylostratigraphic analysis suggests that the emergence of tumour suppressors and oncogenes coincided with the emergence of metazoans (Domazet-Lošo & Tautz, 2010). Cancer susceptibility varies greatly between species (Boddy, Abegglen, et al., 2020; Madsen et al., 2017; Vincze et al., 2022). As mutations arise from imperfect cell replication, larger animals with greater lifespans who undergo a greater number of cell divisions should theoretically be at a higher risk of mutations and consequently cancer. Yet many large-bodied mammals, such as whales (Tollis et al., 2019) and elephants (Abegglen et al., 2015), have extremely low rates of cancer. This lack of correlation between body size and cancer risk is known as Peto’s paradox (Nunney et al., 2015) and has been supported by a number of studies (Abegglen et al., 2015; Boddy, Abegglen, et al., 2020; Bulls et al., 2022; Compton et al., 2025; Vincze et al., 2022). A more recent model of cancer mortality contradicts Peto’s paradox (Butler et al 2025), highlighting that larger mammals often have higher rates of malignancy. However, this relationship was logarithmic, meaning observed cancer rates were still lower than expected relative to body size, suggesting larger mammals may have evolved more efficient anticancer mechanisms.

This differential cancer susceptibility is often attributed to life history trade-offs and antagonistic pleiotropy (Boddy, Harrison, et al., 2020). Life history theory predicts that there is a trade-off between reproduction and maintenance (Kirkwood, 1977). Organisms that allocate more resources to reproduction have less to dedicate to somatic maintenance, which renders them more susceptible to aging-related degeneration such as cancer. This appears to be supported by studies that have shown mammals with large litter sizes and long lactation periods are at greater risk of cancer (Dujon et al., 2023). Antagonistic pleiotropy refers to genes that are ultimately deleterious but confer an advantage earlier in life (Ljubuncic & Reznick, 2009). An example is the *Xmrk* oncogene in *Xiphophorus* fish. Although it causes melanoma (Wittbrodt et al., 1989), *Xmrk* is also associated with traits that increase reproductive success such as a larger size (Fernandez & Bowser, 2010) and a spotted caudal melanin pattern (Fernandez & Morris, 2008).

Comparative genomic studies have shown that there is often a genetic basis for cancer resistance. Some of the most well studied oncogenes are the three canonical Ras genes (*H-Ras, K-Ras* and *N-Ras*). These are responsible for 15-20 % of human cancers (Quinlan & Settleman, 2009) and only require a single mutation at codon 12, 16 or 31 to become activated (Prior et al., 2012). First discovered in the 1960s (Harvey, 1964), *H-Ras, K-Ras* and *N-Ras* were the first identified members of the much larger Ras superfamily, with over 150 genes found across all forms of eukaryotic life (Goitre et al., 2014). The Ras superfamily is divided into five major gene subfamilies: Ras (which includes the three canonical genes), Rho, Rab, Ran and Arf (Goitre et al., 2014).

Duplication of tumour suppressors has also been linked to cancer resistance. For example, the microbat (*Myotis lucifugus*) has 63 copies of *FBXO31* (Caulin et al., 2015). This is a tumour suppressor gene that induces cell cycle arrest in response to DNA damage (Santra et al., 2009). Bats also express high levels of the transporter *ABCB1*, which provides protection against genotoxic agents (Koh et al., 2019). Similar expansions have been observed in the African elephant (*Loxodonta Africana)*, which has 20 copies of the tumour suppressor gene *TP53* (Abegglen et al., 2015). The TP53 pathway is activated in the presence of genotoxic agents and, interestingly, in elephants is activated at lower doses of these agents than other species (Sulak et al., 2016). The bowhead whale (*Balaena mysticetus*), one of the longest living mammals, also has duplications of *PCNA* and species-specific mutations in *ERCC1*, both genes involved in DNA repair (Keane et al., 2015). The famously cancer-resistant naked mole rat (*Heterocephalus glaber*) has higher copy numbers of *TINF2* and *CEBPG*, two genes involved in telomere protection and DNA repair respectively (MacRae et al., 2015), and is documented to have better base and nucleotide excision repair systems than other rodents (Evdokimov et al., 2018).

However, the inverse has rarely been explored – why are some taxa particularly susceptible to cancer? One example of taxa with high cancer susceptibility is the Dasyuridae, a family of small to medium sized marsupial carnivores (García-Navas et al., 2020). Like other marsupials, they give birth to altricial neonates who complete development in the pouch (Old & Deane, 2000). Many dasyurid species give birth to supernumerary young (more offspring than can be supported by the number of available teats) (Gemmell et al., 2002; Parrott & Edwards, 2023). Cancer has been reported in many species of dasyurids (Anderson et al., 1990; Attwood & Woolley, 1973; Hopkins & Gaynor, 1985; Straube & Callinan, 1980; Twin & Pearse, 1986), with the family frequently being referred to as having a particularly high susceptibility to cancer (Attwood & Woolley, 1973; Canfield et al., 1990b; Old & Stannard, 2020; Twin & Pearse, 1986; Vincze et al., 2022). Vincze et al. (2022) observed that the kowari (*Dasyuroides byrnei*) had the highest cancer mortality risk out of the 191 mammalian species included in their comparative study – a value so high that the species was excluded due to concerns that its high leverage would bias the model. Not only is cancer the most common cause of mortality for Tasmanian devils (*Sarcophilus harrisii*) in captivity (Peck et al., 2019), but they are one of only two vertebrates to suffer from contagious cancers (devil facial tumour disease, DFTD), and the only vertebrate to suffer from two separate contagious cancers (DFT1 and DFT2) (Metzger & Goff, 2016). However, it is unknown whether this high incidence of cancer amongst dasyurids arises from a genetic predisposition to cancer, or that they are frequently housed in captivity where artificial conditions and veterinary care lead to increased lifespans compared to individuals in the wild (Jackson, 2007; Obendorf, 1993). It is therefore possible that cancer is simply reported more often in these genera, rather than occurring more often. Another marsupial species, the koala (*Phascolarctos cinereus*), has also shown elevated rates of lymphoma and leukaemia, although this is often linked to koala retrovirus (KoRV) (McEwen et al., 2021; Tarlinton et al., 2005). Retroviruses such as KoRV are known to cause cancer in various species (Hartmann, 2012; Ruprecht et al., 2008; Tarlinton & Greenwood, 2024).

The increasing availability of scaffolded or chromosome level reference genomes has enabled a range of comparative evolutionary studies, including for marsupials, with high quality reference genomes currently available for 18 species (Challis et al., 2023). In order to conduct a more comprehensive comparative study across the marsupial family tree, we generated a long read scaffolded reference genome for the kowari (highest reported marsupial cancer susceptibility) (Vincze et al., 2022) and eastern barred bandicoot (*Perameles gunnii*). Here we aimed to use these genomes, in conjunction with nine other high quality marsupial genomes, to investigate the evolution of cancer related genes in marsupials using a comparative framework.

## Results and Discussion

### Genome Assemblies

We sequenced the kowari and eastern barred bandicoot genome with PacBio HiFi and scaffolded using Hi-C, with the resulting assemblies 3.21Gb and 3.94Gb in size, containing 851 and 1,462 scaffolds for the kowari and eastern barred bandicoot, respectively. In addition, we downloaded nine publicly available genomes (accession numbers and genome information, Table S1). Benchmarking universal single-copy orthologs (BUSCO) analysis revealed highly complete assemblies for all 11 genomes used in this study, with over 89.5% (ten above 94.7%) of single copy mammalian genes identified (Table S1). For more information on the genome assemblies, see Supplementary Material.

### Cancer Prevalence

We collated data on reports of neoplasia in marsupials from five published sources (Canfield et al., 1990a, 1990b; Effron et al., 1977; Ladds, 2009; Ratcliffe, 1933) and the Australian Wildlife Health Information System (eWHIS) (Table S2 & S3). Dasyurids had the highest number of reported neoplasia amongst marsupials, comprising 37.35% (282/757) of all reports. This was even though eWHIS may not include all cases of neoplasia in captive Tasmanian devils (Cox-Witton, K., *pers. comm.*). Within dasyurids, the highest reports of neoplasia occurred in the Tasmanian devil, three quoll species (*Dasyurus hallucatus*, *Dasyurus maculatus*, *Dasyurus viverrinus*) and kowari (Table S3). Koalas had the second highest rates amongst marsupials (27.6%, 209/757), although most of these were from the most recent source (eWHIS, 2008-2024). Nearly all reports (105 out of 113) were koalas from Queensland and New South Wales, which may reflect the higher prevalence of KoRV in these northern regions (Simmons et al., 2012). For this reason, cancer genes were investigated in both dasyurid and koala ancestors.

### Gene Family Evolution

First, we annotated 11 marsupial genomes using FGENESH++v7.2.2 (Solovyev et al., 2006), which annotated between 28,365 and 76,963 genes in all 11 species (Table S4). We then identified orthologs of all genes with mutations implicated in human cancer (*n* = 753), from the Cosmic Cancer Gene Census v101 GRCh38 (Sondka et al., 2024) using a reciprocal blast best hit analysis. We identified between 577 and 688 orthologs of these cancer related genes in each marsupial genome (Table S4).

Genes were also assigned to orthogroups (sets of genes descended from a single ancestral gene) using Orthofinder v2.4.0. In total, 260,261 annotated genes (84.3%) were assigned to 24,416 orthogroups amongst the 11 species. 7,640 (31.29%) of these orthogroups had all species present and 2,999 (12.28%) were single copy orthologs. The opossum had the lowest percentage of genes (68.8 %) in orthogroups, which is expected as it was the phylogenetic outgroup (Table S4).

We undertook a Computational Analysis of gene Family Evolution (CAFE) (Mendes et al., 2020) analysis to identify rapidly evolving orthogroups across marsupial lineages. Across the five marsupial orders (Diprotodontia, Dasyuromorphia, Peramelemorphia, Microbiotheria, Didelphimorphia), CAFE identified 229 orthogroups that had undergone statistically significant expansions and 115 that had undergone statistically significant contractions (Figure 1A). The orthogroups in the koala and dasyurid ancestors were then examined in more detail because of the high cancer prevalence in these two lineages.

**Figure 1.**
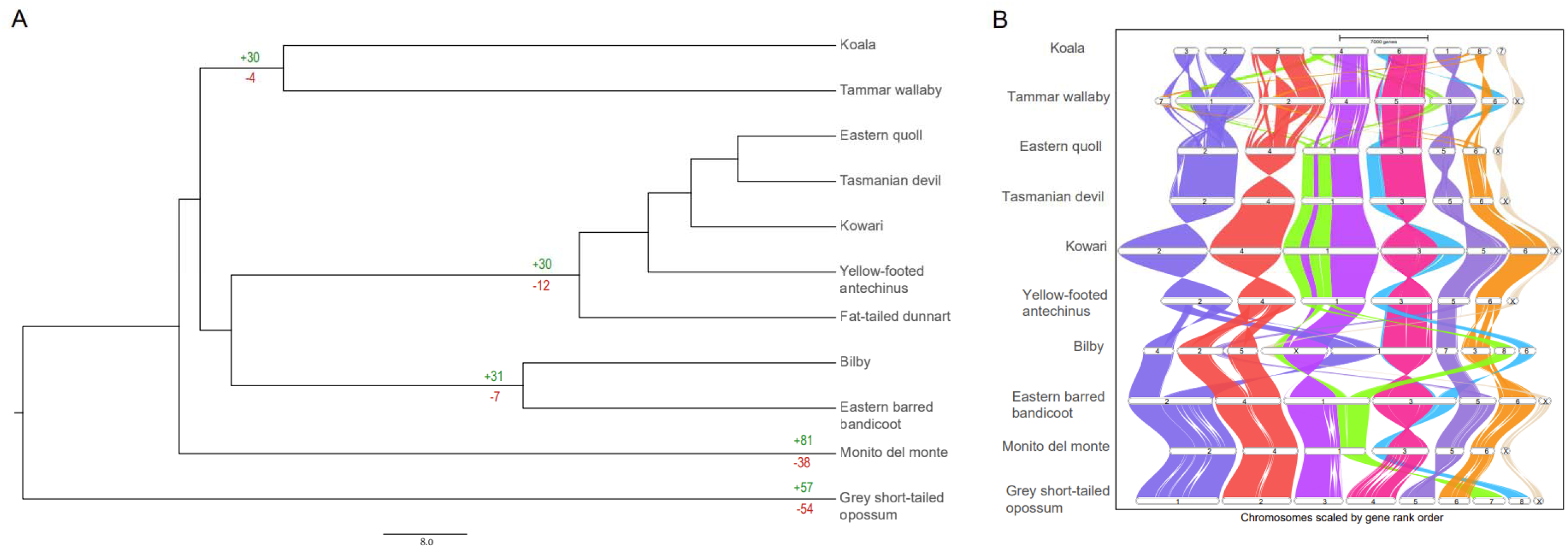
A. For each marsupial order, the number in green above the branch represents the number of orthogroups deemed to have undergone a statistically significant expansion by CAFE and the number in red beneath the branch represents the number of orthogroups deemed to have undergone a statistically significant contraction. The figure was created by manually annotating CAFE output onto the time calibrated phylogeny (see methods for more detail about how phylogenetic tree was generated). The scale bar represents 8 million years. B. Synteny plots generated by GENESPACE. The coloured blocks represent chromosome scaffolds for each species. The opossum was chosen to represent chromosome order as a model for the ancestral species. The dunnart is not included in this plot as the genome was fragmented.

CAFE identified 41 significant expansions and 10 significant contractions in the koala ancestor. However, only five gene families contained cancer related genes from the COSMIC database and all were from the major immune gene families T cell receptors (TCRs) and immunoglobulins (IGs). Two of the signficant gene families in the dasyurid ancestor also involved TCRs and IGs (see below).

Automated pipelines, such as FGENESH++, are known to be unreliable at annotating immune genes, especially those that are highly polymorphic and rapidly evolving such as the TCR and IG variable regions (Peel et al., 2022). The major immune gene families have also been well characterised and compared in marsupials and the constant region gene numbers are conserved. Hogg et al. (2024) identified similar numbers of IG and TCR constant genes in the bilby, devil, short-tailed opossum, koala and woylie (ranging from 11-20 for IGs and 9-14 for TCRs). Previous studies have indicated the koala did have a higher number of IG variable region genes (289 compared to 226 in the woylie and 116 in the bilby), although this high number is attributed to genome quality rather than gene expansion (Peel et al., 2022). Further, gene expansions and contractions are how immune gene families evolve. As immune genes have a basic biological function, it is difficult to disentangle this from their involvement in cancer. For these reasons, orthogroups containing IGs and TCRs were not investigated further.

In the dasyurid ancestor, CAFE identified 30 significant expansions and 12 significant contractions. Only six of these gene families contained cancer related genes from the COSMIC database, and two of these involved TCRs and IGs and so were not investigated further. Three of the remaining four orthogroups underwent contractions in subsequent lineages. For example, orthogroup HOG000767 contained putative orthologs of *ATRX*, a tumour suppressor. Although this underwent a significant expansion in the dasyurid ancestor, it subsequently underwent contractions in the ancestor to the quoll and devil (Figure S3). Orthogroup HOG0001427 contained putative orthologs of *SLC34A2*, a tumour suppressor also involved in oncogenic fusion. This underwent contractions in the quoll and kowari (Figure S4). Orthogroup HOG0002169 contained orthologs of *TCEA1*, a gene involved in oncogenic fusions and subsequently underwent contraction in the devil (Figure S5). As these orthogroups were not consistently high in all dasyurid species, they were not investigated further.

The remaining orthogroup containing cancer genes (HOG0000752) underwent significant expansion in the dasyurid ancestor and had much higher numbers of genes in all extant dasyurids (average 12 ± 4.3 standard deviation) compared to the other marsupials in this study (average 0.67 ± 1.2) (Figure 2). In this context, statistical significance was determined by the CAFE analysis. This orthogroup contained genes annotated as *K-ras*, a well-known oncogene in the Ras gene family.

**Figure 2.**
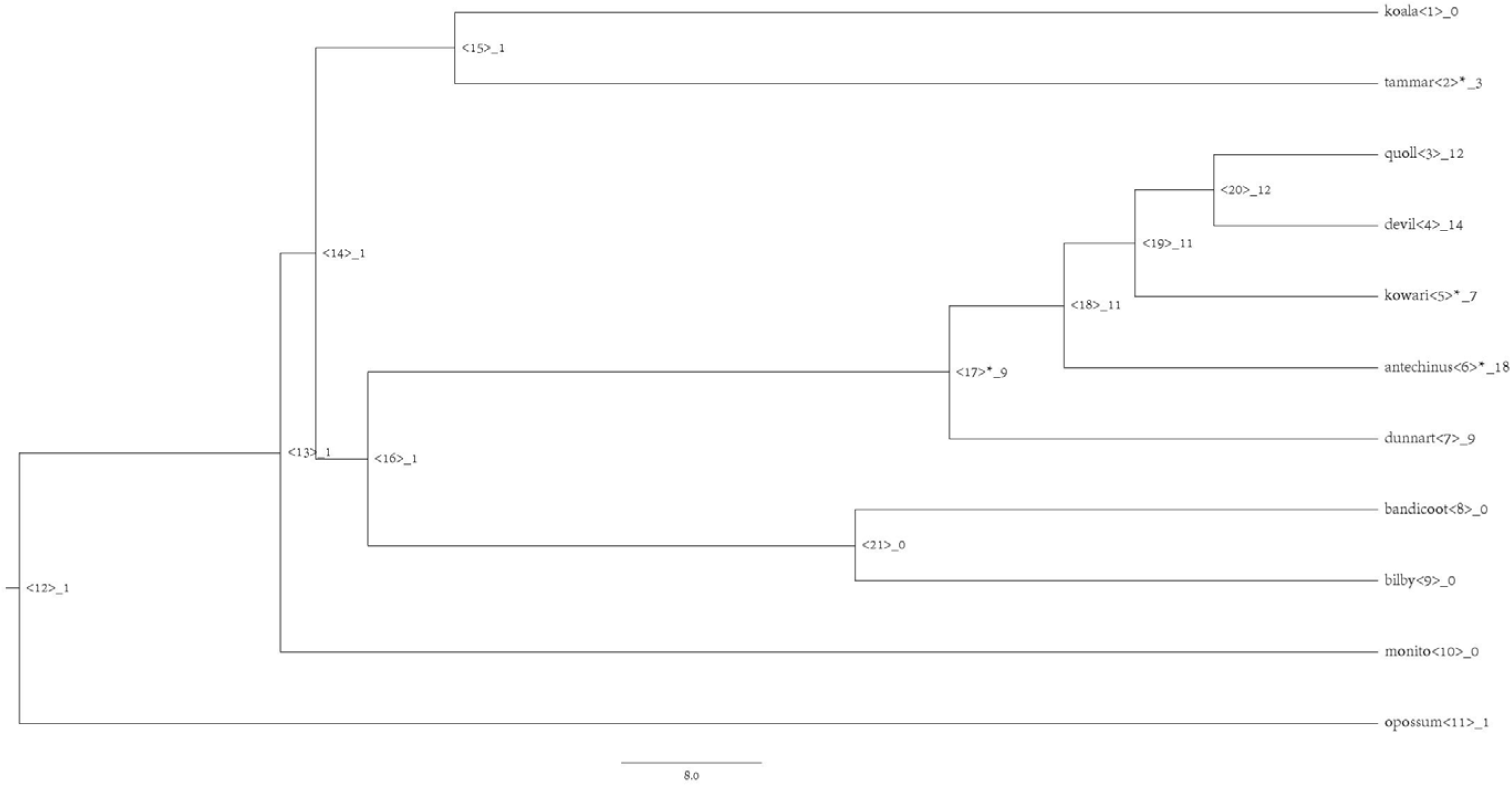
Gene tree for orthogroup HOG0000752 annotated as *K-ras*. The number after the underscore represents the number of genes within the orthogroup encoded in the genome of each species (or is predicted to encode, in the case of ancestral nodes). Asterisk indicates that a statistically significant expansion or contraction occurred in this lineage, as seen in the dasyurid ancestor.

*K-ras* is highly conserved across all jawed vertebrate species and is usually present in single copy within the genome and transcribed in two isoforms (*K-Ras4A* and *K-Ras4B*). All dasyurids and tammar wallaby were predicted to encode between three and 18 copies of genes within this orthogroup (Figure 2). In addition to containing more than one copy, the marsupial genes in this orthogroup showed much higher differentiation compared to *K-ras* from other mammals, birds and amphibians (Figure 3). The amino acid sequence similarity amongst non-marsupials ranged 94.15% (between the African clawed frog and zebra fish) to 100% (between the human and chicken). The sequence similarity amongst marsupials ranged from 68.75% (between the dunnart and the quoll) to 97.12% (between the devil and the kowari). The similarity between marsupials and non-marsupials ranged from 58.54% (between the dunnart the African clawed frog) to 68.75% (between the antechinus and the human, chicken and zebrafish).

**Figure 3.**
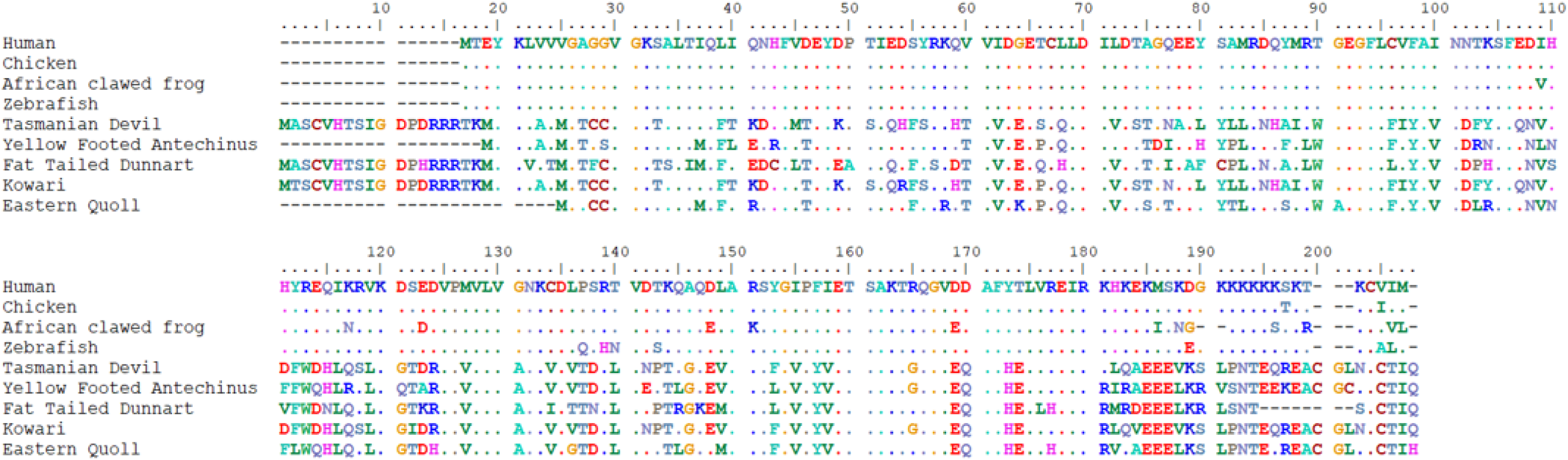
Alignment of a subset of marsupial genes annotated as *K-ras* by FGENESH++ with K-ras genes from eutherian, amphibian, avian and fish species: humans (*Homo sapiens,* NP_001356716.1), chicken (*Gallus gallus,* NP_001243091.1, African clawed frog (*Xenopus laevis,* NP_001081316.1) and zebrafish (*Danio rerio,* NP_001003744.1). The marsupial species are the Tasmanian devil, yellow footed antechinus, fat-tailed dunnart, kowari and eastern quoll. Dots represent 100% amino acid identity to human. All sequences showed high similarity (> 58.6% similarity using the BLOSUM62 similarity matrix) but there were distinct differences between the marsupial sequences and those from other taxa. Only a subset of genes is shown for ease of viewing, as the orthogroup contained 60 dasyurid genes.

### Novel Ras Genes Discovered in Marsupials Annotation

As CAFE identified expansions in the *K-Ras* gene family in the dasyurid ancestor, we chose to further investigate other members of the Ras subfamily. First, to check if the automated annotations were correct, we manually annotated the four canonical Ras genes (*HRas, NRas, KRas4A* and *KRas4B*) in all 11 study species. A single copy of *HRas, NRas, KRas4A* and *KRas4B* were identified in all marsupial species in this study (Supplementary File 1). In addition to these, an almost exact duplicate of *Kras4B* was also annotated in the opossum, with only a single amino acid change (K>Q at 101). This duplicate was encoded by a single exon as opposed to the four exons in the other *K-ras* genes. However, the marsupial genes that FGENESH++ had annotated as *K-Ras* were not amongst these canonical genes, indicating they may be novel members of the Ras gene family.

To confirm that these genes were novel and not orthologs of other members of the Ras gene subfamily, we chose to manually annotate the remaining 33 genes in the Ras subfamily in the Tasmanian devil (the Ras subfamily being one of the five subfamilies that comprise the Ras superfamily). We selected the devil as a representative marsupial as it had the most complete genome amongst the dasyurids. Out of these 33 other genes in Ras subfamily, 29 were identified in the devil (Supplementary File 1), the four we were unable to identify were *ERas, RERGL, RHEBL1* and *DIRAS3*. *ERas* is an unusual gene within the Ras subfamily as it is only encoded by a single exon, the only Ras gene without paralogs and has so far only been characterised in eutherian mammals (De Falco et al., 2022; Roperto et al., 2017; Takahashi et al., 2003; Tanaka et al., 2009). Our results provide further support that *Eras* is not only mammal specific but eutherian specific.

In addition to these genes, we identified a distinct phylogenetic clade of putative novel marsupial *Ras* genes that belonged to the Ras subfamily but did not show orthology with any of the other vertebrate *Ras* genes (Figure 4). Some of these had been incorrectly annotated by FGENESH++ as one of the classical Ras genes (*H-Ras, K-Ras* or *N-Ras*). The genes in this clade were all encoded by a single exon and ranged in length from 155 to 390 amino acids. All putative novel Ras genes contained an open reading frame, a RAS smart domain and at least partial matches for four out of the five conserved G box motifs characteristic of the Ras superfamily (Figure S6). The top reciprocal BLAST hit for all genes was a canonical Ras gene, yet the novel genes contained a more diverse sequence compared to eutherian sequences (Figure 3). These genes were almost exclusively expressed in marsupial gonads (Table 1, Supplementary File 2) and so were named in the order in which they were found using the first letters of the genus and species then MgRas (for marsupial gonad Ras gene).

**Figure 4.**
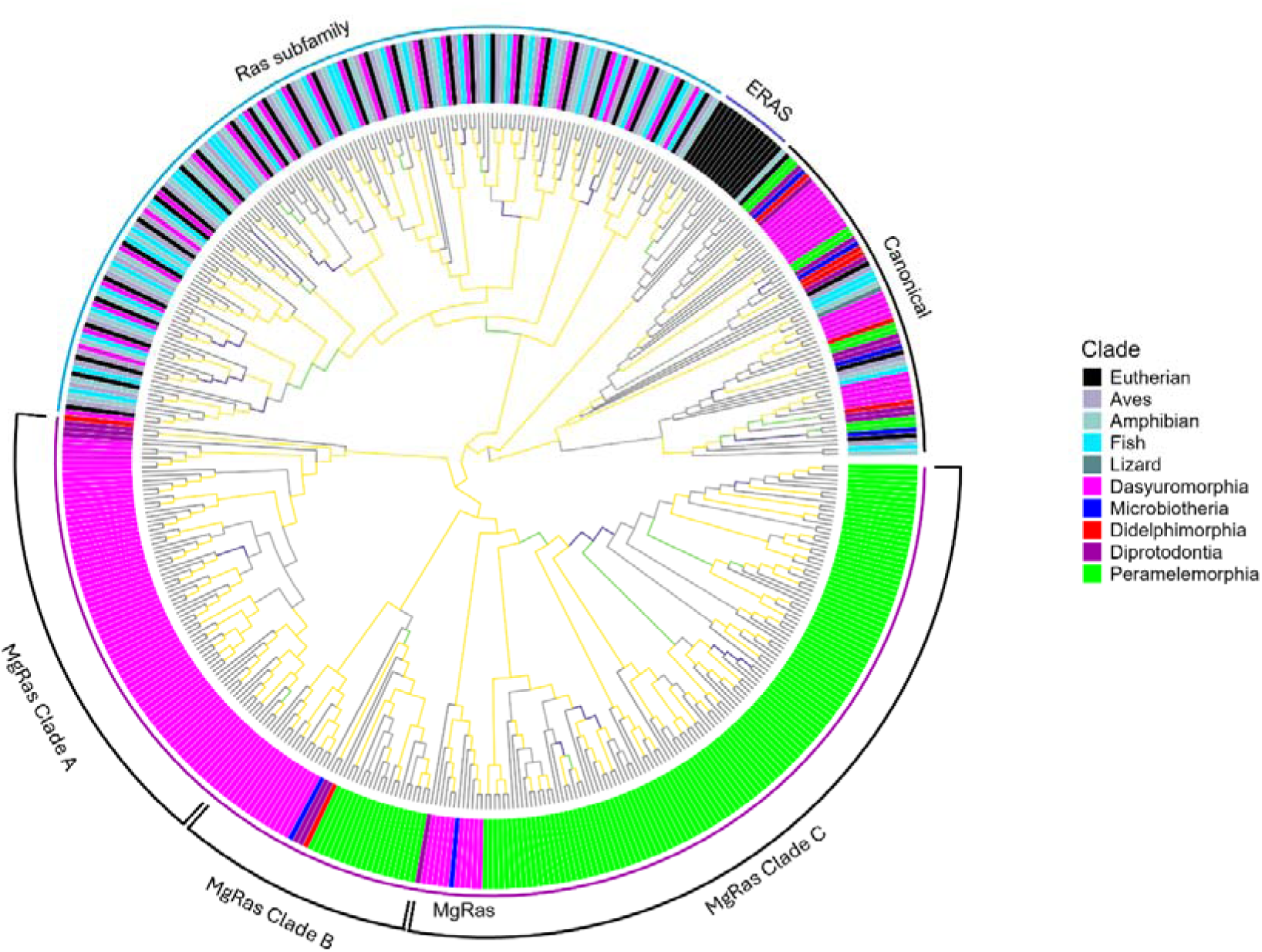
Phylogenetic relationships among Ras subfamily genes in marsupials, eutherians, birds, amphibians, and fish. ERas genes were eutherian specific and form a monophyletic clade, MgRas genes were marsupial specific and formed a monophyletic clade with three subclades (Clade A, Clade B, Clade C). The rest of the Ras subfamily were found in all vertebrate species. Yellow branches indicate bootstrap values between 95 and 100, green branches indicate bootstrap values 90-94, dark blue branches indicate bootstrap values between 80 and 89, grey branches indicate bootstrap values under 80. Terminal branches are in black.

**Table 1.** Number of MgRas genes identified amongst the 11 marsupials in this study and their tissue expression. The first number represents the total number of genes, followed by how many genes belonged to each phylogenetic clade (A, B and C). N/A indicates that gonad transcriptome was not available for that species. NCBI accession numbers for genomes and RNAseq data are included in Table S1. For the opossum, additional RNAseq data from PRJNA193216 was used (Wang et al., 2014).

| Order | Species | Number of genes | Number of genes expressed in gonads | Number of genes expressed in other tissues |
| --- | --- | --- | --- | --- |
| Diprotodontia | Koala | 3<br>(A: 1, B: 1, C: 1) | 3 | 1 (pouch) |
|  | Tammar wallaby | 3<br>(A: 2, B: 0, C: 1) | N/A | 0 |
| Dasyuromorphia | Eastern quoll | 22<br>(A: 16, B: 2, C: 4) | N/A | 0 |
|  | Tasmanian devil | 16<br>(A: 14, B: 1, C: 1) | 15 | 0 |
|  | Kowari | 21<br>(A: 15, B: 2, C: 4) | N/A | 1 (spleen) |
|  | Yellow footed antechinus | 23<br>(A: 17, B: 2, C: 4) | 4 | 1 (liver) |
|  | Fat tailed dunnart | 14<br>(A: 10, B: 2, C: 2) | 10 | 1 (proximal yolk sac, distal yolk sac) |
| Peramelemorphia | Greater bilby | 23<br>(A: 0, B: 17, C: 6) | 14 | 0 |
|  | Eastern barred bandicoot | 117<br>(A: 0, B: 103, C: 14) | 77 | 1 (brain) |
| Microbiotheria | Monito del Monte | 2<br>(A: 0, B: 1, C: 1) | N/A | 0 |
| Didelphimorphia | Grey short-tailed opossum | 2<br>(A: 1, B: 0, C: 1) | N/A | 1 (extra embryo membrane, embryo brain) |

Phylogenetic analysis revealed strong support for the MgRas genes being within the Ras subfamily of the Ras superfamily (Figure 4). The MgRas genes and the majority of non-canonical Ras genes each form a monophyletic clade with strong bootstrap support (100% bootstrap) and form a single clade (98% bootstrap support) that sits sister to ERas genes (100% bootstrap). This combined group is, in turn, sister to the classical Ras genes *H-Ras*, *K-Ras* and *N-Ras* (100% bootstrap). Within each clade (canonical, ERas, MgRas, and non-canonical), genes cluster in orthologous groups amongst species, particularly evident for Ras subfamily and canonical Ras clades. Although, within the MgRas clade, distinct order-specific expansions were also evident in the Dasyuromorphia (antechinus, devil, dunnart, kowari and quoll) and the Peramelemorphia (bandicoot and bilby), with all other marsupial species having either two or three MgRas orthologs (Figure 4).

### Genomic Organisation and Synteny

There was strong support (>98% bootstrap) for three phylogenetic clades (termed A, B and C) (Figure 4) that were largely encoded in two main clusters in the genome of each species (Figure 5), as is common in genes that evolve by tandem duplication (Pan & Zhang, 2008). Genes from all three clades showed similar expression patterns.

**Figure 5.**
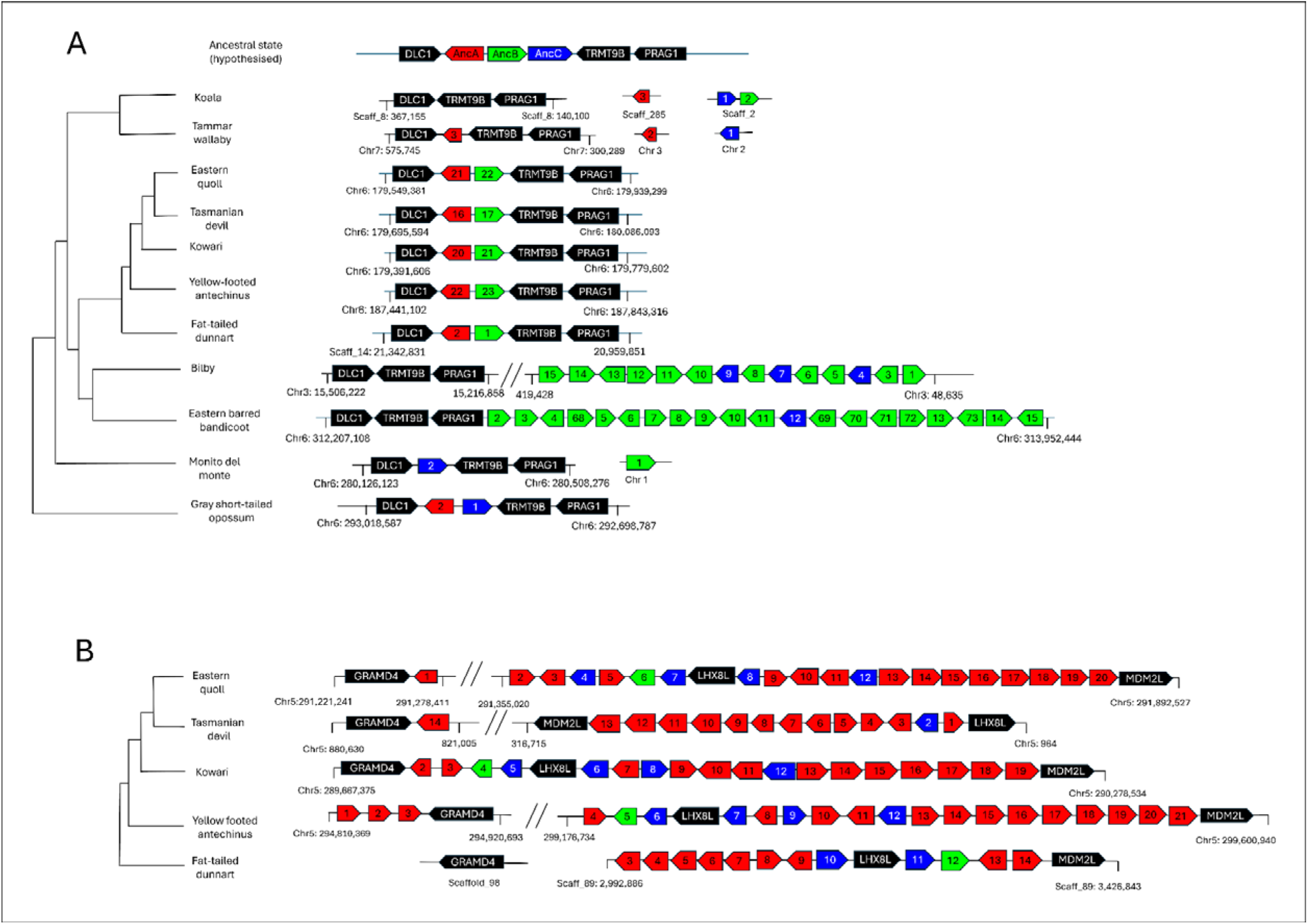
MgRas genes occurred in two main clusters. **(A)** Cluster 1 was found in all marsupials on syntenic chromosomes. Not all bandicoot genes are depicted in figure as the cluster contained 115 genes. The hypothesised ancestral state depicted represents the most parsimonious scenario. **(B)** Cluster 2 was found only in dasyurids. We hypothesise that the ancestral three gene cluster was duplicated and translocated to chromosome 5 in the Dasyuridae. MgRas genes are coloured according to clade (red for A, green for B, blue for C) and flanking genes are in black. The numbers represent the gene number given to each gene.

Cluster 1 was encoded in syntenic chromosomes in all species, although in the diprotodonts (koala and tammar wallaby) and monito del monte a small number of orphan genes were identified on different chromosomes. Cluster 1 contained representatives from all phylogenetic clades, although not all clades were present in all species (Fig 5A). Genes from Clade A were not identified in the monito del monte and Peramelemorphia, and genes from Clade C were not identified in Cluster 1 in the Dasyuridae. In the Peramelemorphia, Clade B underwent an extremely large expansion and Clade C underwent a smaller expansion, relative to other marsupials.

Cluster 2 was only found in dasyurids on chr 5 (scaffold 89 in the dunnart) (Fig 5B). This cluster also contained representatives from all phylogenetic clades, with Clade A having undergone a large expansion and Clade C having undergone a smaller expansion in some species.

All species had some combination of genes from all three clades either flanked or near *DLC1* (Deleted in Liver Cancer 1) and *TRMT9B* (probable tRNA methyltransferase 9B) genes, so the most parsimonious explanation is that the ancestral state included a three gene cluster in this region, which underwent subsequent expansions and contractions in the different marsupial lineages. We hypothesise the entire three gene cluster was duplicated and translocated to chromosome 5 in the dasyurid lineage, where clades A and C subsequently underwent expansions to form Cluster 2.

Both clusters were encoded near potential tumour suppressors (*DLC1, TRMT9B* and GRAM domain containing 4, *GRAMD4*) (Begley et al., 2013; John et al., 2011; Zhang & Li, 2020), with Cluster 1 entirely flanked by tumour suppressors in most species (Figure 5). Oncogenes and tumour suppressors are often in close proximity within the genome (Antonio & Widegren, 2005), and oncogenes without a neighbouring tumour suppressor (within 1.46 Mbp) are more prone to amplification (Wu et al., 2017). The two clusters both occurred near chromosome ends, which is also common in oncogenes (Antonio & Widegren, 2005; Lima-de-Faria et al., 1991).

### Marsupial Ras Genes and Cancer

All genes in the Ras superfamily are molecular switches that regulate a diverse range of functions, although those in the Ras subfamily are primarily involved in cell proliferation, cell growth and the cell cycle (Goitre et al., 2014). Of the 36 mammalian Ras subfamily genes, five are listed in the COSMIC census as oncogenes (*K-Ras, H-Ras, N-Ras, Rap1B, R-Ras2*), with a number of others also implicated in cancer (Berger et al., 2014; Khalil & Nemer, 2020; Suarez-Cabrera et al., 2021; Thies et al., 2021). We hypothesise that, similar to these members of the Ras subfamily, at least some MgRas genes may also function as oncogenes. The expansion observed in dasyurids may render the species more susceptible to cancer, as a greater number of genes increases the probability of incurring a mutation in one.

Interestingly, some members of the Ras subfamily (eg. *NKIRAS1*, *DIRAS3)* appear to have the opposite role and function as tumour suppressors (Bildik et al., 2022; Postler et al., 2023). MgRas gene expansions were also observed in bandicoots and bilbies, two species that had low reported rates of cancer but noting both species have short lifespans (Lynch, 2008). As the major expansion in the Peramelemorphia was not orthologous to the major expansion in the dasyurids (Figure 4) it is possible the different groups of genes have evolved different functions. We recommend future studies use marsupial cancer biopsies to determine (i) if these genes are expressed, and (ii) if these genes (or regulatory regions) have been mutated in cancer tissue.

Oncogenes are retained as they play vital roles in organism function (Spandidos & Anderson, 1989). As the MgRas genes were almost exclusively expressed in the gonads, we hypothesise that their role may be in reproduction. The expansions observed in the Dasyuromorphia and Peramelemorphia may be due to the unique reproductive biology of these two orders. The Dasyuromorphia species in our study all give birth to supernumerary young – an excess number of offspring than the number of teats that can support them (Guiler, 1970; Morton & Fletcher, 1989; Woolley, 1971). Increased litter size has frequently been hypothesized to contribute to cancer risk in species due to antagonistic pleiotropy (Boddy et al 2020), and studies have found that cancer is more likely to be detected in species with larger litter sizes (Dujon et al., 2023)

The Peramelemorphia also have a unique reproductive biology in that they are the only marsupial species to have a chorioallantoic placenta (Renfree, 2010; Tyndale-Biscoe & Renfree, 1987). Although this is more energetically demanding pregnancy, more invasive placentation is interestingly not correlated with increased cancer risk (Dujon et al., 2023). Ultimately future studies should use transcriptomes from reproductive organs taken at different times to determine if gene expression changes throughout reproduction, noting that the species in this order are all currently listed as threatened species so sampling will be difficult.

## Conclusion

Here, we aimed to characterise the evolution of cancer related genes in marsupials to determine if cancer susceptibility in certain lineages may have a genetic basis. Our results show a novel marsupial specific lineage of Ras genes that has undergone unique family specific expansions in both the Dasyuromorphia and the Peramelemorphia species (bandicoot and bilby). As these genes are expressed in the reproductive organs, we theorise that they have expanded as a result of the supernumery offspring in the Daysuromorphia, and invasive placentation in the Peramelemorphia. Genomic organisation suggests some of these may function as oncogenes with the potential to influence cancer mortality risk in some species and recommend future studies to verify gene function.

## Materials and Methods

### Eastern Barred Bandicoot and Kowari Reference Genome Assemblies

Three frozen kowari tissue samples were obtained from the Australian Biological Tissue Collection (ABTC, South Australian Museum): liver from one individual (Reg No ABTC7551) and heart and spleen from a second individual (Reg No ABTC7553). Three bandicoot tissues samples were obtained from two individuals: kidney from a female (University of Melbourne); and brain and gonad from a male. Both bandicoots were wild individuals from Victoria (the female was found deceased at Mount Rothwell, the male was euthanized after a vehicle strike on Philip Island). The kidney tissue was flash frozen and stored at -80° C. The brain and gonad tissue were stored at -80° C.

High molecular weight DNA was extracted from the kowari heart and bandicoot kidney tissues using the Circulomics Nanobind Tissue Big DNA Kit (Circulomics) (NB-900-701-001) and DNA concentration was measured using Qubit fluorometer (ThermoFisher Scientific). DNA was then pooled for each species and sent to Australian Genome Research Facility (Brisbane, Australia) for PacBio HiFi library prep and Revio sequencing on a 25M SMRT cell. Heart tissue for the kowari and bandicoot was sent for HiC Armina 2.0 library preparation and sequenced on NovaseqX at Biomolecular Resource Facility (Australian National University).

Total RNA was extracted from all three kowari tissues (liver, heart and spleen) and two bandicoot tissues (brain and gonad) using Qiagen RNeasy mini kit (Qiagen, Cat. No. 74104) with on-column DNAse digestion using the DNase I set (Qiagen). RNA quality was assessed using the RNA nano 6000 kit on the Bioanalyzer (Agilent) then submitted to Ramaciotti Centre for Genomics (The University of New South Wales) for Illumina stranded mRNA prep. The RNA was sequenced as 150 bp paired end reads on a NovaSeq X Plus 10B flowcell, resulting in 76 to 120 million reads per sample.

Both reference genomes were assembled on the Galaxy Australia webserver (https://usegalaxy.org.au/) using the Vertebrate Genome Project genome assembly pipeline (Lariviere et al., 2024). Briefly, HiFi reads were quality trimmed and reads containing adapters removed using Cutadapt v4.9 (Martin, 2011), genome size and k-mers were estimated using Meryl v1.3 (Rhie et al., 2020) and GenomeScope v2.0 (Ranallo-Benavidez et al., 2020). Genome assembly was performed using the hifiasm v2.1 in HiC mode (Cheng et al., 2021; Price, 2022) using HiFi reads to assemble contigs and Hi-C reads to identify haplotypes, with the assembly quality assessed using gfastats v1.3.9 (Formenti et al., 2022) and BUSCO v5.8.0 with mammlia_odb10 lineages (Simão et al., 2015). Hi-C reads were then used to scaffold contigs of the primary assembly, scaffolding was performed using YaHS v1.2a.2 (Zhou et al., 2023) with contact maps generated with Pretext v0.1.9 (Harry).

To identify sex chromosomes in the kowari, HiFi reads were mapped to the assembly using minimap2 v2.22-r1101 (Li, 2018) and secondary alignments excluded using Samtools v1.9 (Danecek et al., 2021). The genomes were divided into 1 Mbp windows and the mean coverage for each window was calculated using Bedtools v2.31.0 (Quinlan & Hall, 2010). We also performed NBLASTN and TBLASTN using Blast v2.2.30 (Altschul et al., 1990) search for the sex-linked SRY gene on the Y chromosome, using the human (NP_003131.1 and NM_003140.3) and devil (XP_031801149.1 and XM_031945289.1) SRY genes as query sequences.

### Genome Annotations

Nine additional marsupial genomes were used in this study: Monito del monte (*Dromiciops gliroides*) (Rhie et al., 2021), gray short-tailed opossum (*Monodelphis domestica*) (Rhie et al., 2021), greater bilby (*Macrotis lagotis*) (Hogg et al., 2024), koala (*Phascolarctos cinereus*) (Damas et al., 2022), tammar wallaby (*Notamacropus eugenii*) (O’Neill, R., *pers. comm*.), fat-tailed dunnart (*Sminthopsis crassicaudata*) (Ibeh et al., 2024), yellow-footed antechinus (*Antechinus flavipes*) (Tian et al., 2022), Tasmanian devil (*Sarcophilus harrisii*) (Stammnitz et al., 2023) and Eastern quoll (*Dasyurus viverrinus*) (Hartley et al., 2024). See Table S1 for further information about these genomes. All eleven genomes were annotated using the same method to ensure consistency and reliability of results. For more details about the methods used for genome annotation, see Supplementary Material. A synteny map was created using GENESPACE v1.3.1(Lovell et al., 2022)

### Cancer Gene Prevalence and Orthologs

To estimate cancer prevalence in different marsupial taxa, we collated data from previous published summaries that contained comparisons of different marsupial species. We also obtained data from the Wildlife Health Information System (eWHIS).

The Cancer Gene Census (v100_GRCh38) was downloaded from the COSMIC (Catalogue of Somatic Mutations in Cancer) website https://cancer.sanger.ac.uk/cosmic (Sondka et al., 2024). We extracted coding sequences for these genes from the human reference genome (GCF_000001405.40_GRCh38.p14_cds_from_genomic.fna).

To identify orthologs of these genes in the marsupial species, we performed a reciprocal blast best hit analysis. We used BLASTP v2.2.30 (Altschul et al., 1990) using the marsupial annotation files as queries against the human annotation file GCF_000001405.40_GRCh38.p14 (Schneider et al., 2017) and an e-value cutoff of 0.003. We then performed the reverse i.e. the human annotation file was used as a query against each marsupial annotation file. Two genes (e.g. *A* and *B*) were then deemed to be orthologs if gene *A* was the best hit for gene *B* in the first blast search, and gene *B* was also the best hit for gene *A* in the second blast search. Orthologs of the genes from the Cancer Gene Census were then flagged for easy identification in downstream analysis.

### Gene Family Analysis

To remove potential pseudogenes, we excluded any predicted genes from the FGENESH++ genome annotations that had no mRNA evidence and did not have any hits with any known proteins. We also excluded any genes annotated as repetitive elements (SINE, LINE, L1TD1, transposase, retroposon, retrotransposon, reverse transcriptase). All eleven filtered genome annotations were used as input for Orthofinder v2.4.0 (Emms & Kelly, 2019) to infer orthogroups (sets of genes descended from a single ancestral gene).

We then created a rooted, ultrametric phylogenetic tree. We used a species tree inferred by Orthofinder (SpeciesTree_rooted_at_outgroup_6) as the topology of the tree matched the current marsupial phylogeny consensus (Duchêne et al., 2018; Westerman et al., 2016). We calibrated this tree using r8s v1.81 (Sanderson, 2003) with divergence dates obtained from timetree.org (Kumar et al., 2022) (Table S5).

Highly variable orthogroups with more than 100 genes in a single lineage were analysed separately, as recommended by the CAFE manual (Mendes et al., 2020). We then ran CAFE5 v5.1.0 (Mendes et al., 2020) to estimate an error model for all future analyses (-e flag). We choose a gamma model with k=2 (meaning gene families can belong to one of two different evolutionary rate categories) with two λ values (λ_1_ for Australian marsupials and λ_2_ for American marsupials). The λ parameter refers to the gene family evolution rate, or the probability that a gene will be gained or lost in a particular lineage. For more information about why this model was chosen, see Supplementary Material. Gene families that had undergone significant expansions or contractions in either the koala or dasyurid ancestral lineages (the two lineages with high cancer prevalence) were then manually examined for interesting patterns.

### Manual Gene Annotation

As the CAFE analysis indicated potential expansion of Ras genes in some lineages, we verified the automated gene annotation through manual annotation. Human sequences for the four canonical RAS genes (*H-Ras, K-Ras4A, Kras4B* and *N-Ras*) were downloaded from Uniprot and used as query sequences for TBLASTN v2.2.30 against all the marsupial genomes and transcriptomes. Putative genes were extracted using Bedtools v2.29.2 (Quinlan & Hall, 2010) then inspected using Integrative Genomics Viewer v2.16.0 (Robinson et al., 2011) and exon boundaries identified using either the GT/AG or GC/AG convention. Genes were confirmed to be orthologs if they exhibited the extremely conserved sequences of canonical Ras genes in other vertebrate (García-España & Philips, 2023).

Once orthologs of the canonical genes had been identified in each marsupial species, a hidden Markov model (HMM) was constructed using the nucleotide sequences for each species. The HMM was then used to search all genomes using HMMR v3.3.2 (Eddy, 2009). For each marsupial species, this process (TBLASTN and HMMR) was repeated iteratively, incorporating new sequences into the queries until no new hits were found.

Putative hits were first checked for an open reading frame (including start codon and stop codon) and aligned using ClustalW in BioEdit (Hall, 1999). They were initially retained if they contained partial sequences for at least four out of the five conserved G box motifs of Ras genes (GXXXXGKS/T, T, DXXG, T/NKXD, C/SAK/L/T) (Colicelli, 2004). For each sequence, BLASTP was then performed against non-redundant sequences in *Homo sapiens* using the webserver https://blast.ncbi.nlm.nih.gov/ (*Homo sapiens* was used as it is well annotated). The SMART webserver (http://smart.embl-heidelberg.de/) (Letunic et al., 2021) was also used to identify domains in the sequences. Putative genes were then considered to be Ras genes if the best BLAST hit was a canonical Ras gene (*H-Ras, K-Ras, N-Ras*) and the SMART domain was also Ras. We used featureCounts in the subread package v1.5.1 (Liao et al., 2014) to determine if these genes were expressed in any of the tissues in the available transcriptomes. Genes were considered to be expressed if they had at least three reads present in a transcriptome.

To determine the position of the newly annotated genes in RAS phylogeny, we also annotated 29 out of the 33 Ras subfamily genes in the devil genome using the same method (see Table S6 for details of the query sequences). We then aligned nucleotide sequences using MAFFT 7.522 (Katoh & Standley, 2013)and generated a phylogenetic tree with IQTree v2.2.2 (Minh et al., 2020) using ModelFinder (Kalyaanamoorthy et al., 2017) to determine the best model. For branch support, we used 1000 replicates of ultrafast bootstrap (Hoang et al., 2018) and an SH-aLRT test (Guindon et al., 2010) with 1000 replicates. The tree was rooted using the three canonical Ras genes (*K-Ras*, *N-Ras* and *H-Ras*) as these are the founding members of the gene family. Accession numbers for genes used to create the tree are included in Table S7.

In order to identify patterns of synteny, we annotated flanking genes around Ras expansions by manually inspecting the devil and antechinus genomes in NCBI’s Genome Data Viewer (Rangwala et al., 2021). We then used these as query sequences in TBLASN against the other genomes.

ChatGPT 3.5 (https://chatgpt.com/) was used for assistance in some coding and debugging, and all scripts were carefully evaluated and remain the responsibility of the authors.

## Supporting information

Supplemental File 1 Marsupial Ras Annotations

Supplemental File 2 Feature counts

## Acknowledgements

We would like to thank Wildlife Health Australia for facilitating provision of data from the national Wildlife Health Information System (eWHIS). Wildlife health data were provided by: Adelaide Koala and Wildlife Centre, Agriculture Victoria, Australia Zoo Wildlife Hospital, Bonorong Wildlife Sanctuary, Currumbin Wildlife Sanctuary, Department of Natural Resources and Environment Tasmania, Department of Primary Industries and Regions SA, Healesville Sanctuary, James Cook University, Kingston Animal Hospital, Melbourne Zoo, NSW Department of Primary Industries, Perth Zoo, Queensland Department of Agriculture and Fisheries, RSPCA Qld, Taronga Western Plains Zoo, Taronga Zoo Wildlife Hospital, University of Queensland, WA Department of Primary Industries and Regional Development, WA Wildlife, and Zoos SA. We would like to acknowledge the contribution of the Threatened Species Initiative Consortium in the generation of data used in this publication. The Initiative is supported by funding from Bioplatforms Australia, enabled by the Commonwealth Government National Collaborative Research Infrastructure Strategy (NCRIS) in partnership with the University of Sydney; the Australian Government Department of Climate Change, Energy, the Environment and Water; WA Department of Biodiversity, Conservation & Attractions; and Amazon Web Services. We would like to thank the South Australian Museum for the kowari samples, A. Weeks and D. Sutherland for the eastern barred bandicoot samples. We thank the Vertebrate Genome Project for the generation of the Monito del Monte genome, with sequencing and genome assembly conducted by the Max Planck Institute in Dresden, led by Gene Meyers, and coordinated with Olivier Fedrigo and Erich D. Jarvis; and the Gray short-tailed opossum genome, with sequencing and genome assembly conducted at the Vertebrate Genomes Lab (VGL) at the Rockefeller University, led by Olivier Fedrigo and Erich D. Jarvis, with support from Ed Lien and Trygve Bakken at the Allen Institute for Brain Science.

## Funding

This work was supported by funding from the ARC Centre of Excellence for Innovations in Peptide and Protein Science (CE200100012) and the NCRIS-funded Bioplatforms Australia Threatened Species Initiative.

## Data Availability

The genome assemblies and raw transcriptome files are available on NCBI and the SRA, under BioProject PRJNA1301011 (kowari) and PRJNA1301027 (Eastern barred bandicoot).

## Author Contributions

CP designed the study and performed comparative genomics analyses with guidance from EP and LWS. CP and LWS generated genome assemblies for the kowari and Eastern barred bandicoot and genome annotations for all species. RJO and PGSG generated the tammar wallaby genome assembly. CJH and KB sourced funding, samples, and undertook project management and supervision. CP wrote the main manuscript text, with feedback and revisions on the manuscript were provided by all authors.

## Notes

### Competing Interest Statement

The authors have declared no competing interest.

## References

Abegglen, L. M., Caulin, A. F., Chan, A., Lee, K., Robinson, R., Campbell, M. S., Kiso, W. K., Schmitt, D. L., Waddell, P. J., & Bhaskara, S. (2015). Potential mechanisms for cancer resistance in elephants and comparative cellular response to DNA damage in humans. Jama, 314(17), 1850–1860. DOI:10.3390/genes14030546

Aktipis, C. A., Boddy, A. M., Jansen, G., Hibner, U., Hochberg, M. E., Maley, C. C., & Wilkinson, G. S. (2015). Cancer across the tree of life: cooperation and cheating in multicellularity. Philosophical Transactions of the Royal Society B: Biological Sciences, 370(1673), 20140219.

Altschul, S. F., Gish, W., Miller, W., Myers, E. W., & Lipman, D. J. (1990). Basic local alignment search tool. Journal of molecular biology, 215(3), 403–410.

Anderson, W. I., Peters, D. N., Stoffregen, D. A., Steinberg, H., & Wallace, C. (1990). Glossal squamous cell carcinoma with pulmonary metastasis in a kowari (Dasyuroides byrnei). Australian veterinary journal, 67(1), 29. 10.1111/j.1751-0813.1990.tb07389.x

Antonio, L.-D.-F., & Widegren, B. (2005). The non-random location of human oncogenes and tumour suppressor genes. Caryologia, 58(1), 1–14.

Arnal, A., Tissot, T., Ujvari, B., Nunney, L., Solary, E., Laplane, L., Bonhomme, F., Vittecoq, M., Tasiemski, A., & Renaud, F. (2016). The guardians of inherited oncogenic vulnerabilities. Evolution, 70(1), 1–6.

Attwood, H., & Woolley, P. (1973). Spontaneous malignant neoplasms in dasyurid marsupials. Journal of Comparative Pathology, 83(4), 569–581.

Begley, U., Sosa, M. S., Avivar-Valderas, A., Patil, A., Endres, L., Estrada, Y., Chan, C. T., Su, D., Dedon, P. C., & Aguirre-Ghiso, J. A. (2013). A human tRNA methyltransferase 9-like protein prevents tumour growth by regulating LIN9 and HIF1-α. EMBO molecular medicine, 5(3), 366–383.

Berger, A., Imielinski, M., Duke, F., Wala, J., Kaplan, N., Shi, G., Andres, D., & Meyerson, M. (2014). Oncogenic RIT1 mutations in lung adenocarcinoma. Oncogene, 33(35), 4418–4423.

Bildik, G., Liang, X., Sutton, M. N., Bast Jr, R. C., & Lu, Z. (2022). DIRAS3: an imprinted tumor suppressor gene that regulates RAS and PI3K-driven cancer growth, motility, autophagy, and tumor dormancy. Molecular cancer therapeutics, 21(1), 25–37.

Boddy, A. M., Abegglen, L. M., Pessier, A. P., Aktipis, A., Schiffman, J. D., Maley, C. C., & Witte, C. (2020). Lifetime cancer prevalence and life history traits in mammals. Evolution, medicine, and public health, 2020(1), 187–195.

Boddy, A. M., Harrison, T. M., & Abegglen, L. M. (2020). Comparative oncology: New insights into an ancient disease. Iscience, 23(8), 101373.

Bulls, S. E., Platner, L., Ayub, W., Moreno, N., Arditi, J.-P., Dreyer, S., McCain, S., Wagner, P., Burgstaller, S., & Davis, L. R. (2022). Cancer prevalence is related to body mass and lifespan in tetrapods and remarkably low in turtles. bioRxiv, 2022.2007. 2012.499088.

Canfield, P., Hartley, W., & Reddacliff, G. (1990a). Spontaneous proliferations in Australian marsupials—a survey and review. 1. Macropods, koalas, wombats, possums and gliders. Journal of Comparative Pathology, 103(2), 135–146.

Canfield, P., Hartley, W., & Reddacliff, G. (1990b). Spontaneous proliferations in Australian marsupials—a survey and review. 2. Dasyurids and bandicoots. Journal of Comparative Pathology, 103(2), 147–158.

Caulin, A. F., Graham, T. A., Wang, L.-S., & Maley, C. C. (2015). Solutions to Peto’s paradox revealed by mathematical modelling and cross-species cancer gene analysis. Philosophical Transactions of the Royal Society B: Biological Sciences, 370(1673), 20140222.

Challis, R., Kumar, S., Sotero-Caio, C., Brown, M., & Blaxter, M. (2023). Genomes on a Tree (GoaT): a versatile, scalable search engine for genomic and sequencing project metadata across the eukaryotic tree of life. Wellcome open research, 8.

Chan, R. J., & Feng, G.-S. (2007). PTPN11 is the first identified proto-oncogene that encodes a tyrosine phosphatase. Blood, 109(3), 862–867.

Cheng, H., Concepcion, G. T., Feng, X., Zhang, H., & Li, H. (2021). Haplotype-resolved de novo assembly using phased assembly graphs with hifiasm. Nature methods, 18(2), 170–175.

Colicelli, J. (2004). Human RAS superfamily proteins and related GTPases. Science’s STKE, 2004(250), re13–re13.

Compton, Z. T., Mellon, W., Harris, V. K., Rupp, S., Mallo, D., Kapsetaki, S. E., Wilmot, M., Kennington, R., Noble, K., & Baciu, C. (2025). Cancer prevalence across vertebrates. Cancer discovery, 15(1), 227–244.

Damas, J., Corbo, M., Kim, J., Turner-Maier, J., Farré, M., Larkin, D. M., Ryder, O. A., Steiner, C., Houck, M. L., & Hall, S. (2022). Evolution of the ancestral mammalian karyotype and syntenic regions. Proceedings of the National Academy of Sciences, 119(40), e2209139119.

Danecek, P., Bonfield, J. K., Liddle, J., Marshall, J., Ohan, V., Pollard, M. O., Whitwham, A., Keane, T., McCarthy, S. A., & Davies, R. M. (2021). Twelve years of SAMtools and BCFtools. GigaScience, 10(2), giab008.

De Falco, F., Perillo, A., Del Piero, F., Del Prete, C., Zizzo, N., Marcus, I., & Roperto, S. (2022). ERAS is constitutively expressed in the tissues of adult horses and may be a key player in basal autophagy. Frontiers in veterinary science, 9, 818294.

Domazet-Lošo, T., & Tautz, D. (2010). Phylostratigraphic tracking of cancer genes suggests a link to the emergence of multicellularity in metazoa. BMC biology, 8(1), 1–10.

Duchêne, D. A., Bragg, J. G., Duchêne, S., Neaves, L. E., Potter, S., Moritz, C., Johnson, R. N., Ho, S. Y., & Eldridge, M. D. (2018). Analysis of phylogenomic tree space resolves relationships among marsupial families. Systematic Biology, 67(3), 400–412.

Dujon, A. M., Vincze, O., Lemaitre, J.-F., Alix-Panabières, C., Pujol, P., Giraudeau, M., Ujvari, B., & Thomas, F. (2023). The effect of placentation type, litter size, lactation and gestation length on cancer risk in mammals. Proceedings of the Royal Society B, 290(2001), 20230940.

Eddy, S. R. (2009). A new generation of homology search tools based on probabilistic inference. In Genome Informatics 2009: Genome Informatics Series Vol. 23 (pp. 205–211). World Scientific.

Effron, M., Griner, L., & Benirschke, K. (1977). Nature and rate of neoplasia found in captive wild mammals, birds, and reptiles at necropsy. Journal of the National Cancer Institute, 59(1), 185–198.

Emms, D. M., & Kelly, S. (2019). OrthoFinder: phylogenetic orthology inference for comparative genomics. Genome biology, 20, 1–14.

Evdokimov, A., Kutuzov, M., Petruseva, I., Lukjanchikova, N., Kashina, E., Kolova, E., Zemerova, T., Romanenko, S., Perelman, P., & Prokopov, D. (2018). Naked mole rat cells display more efficient excision repair than mouse cells. Aging (Albany NY), 10(6), 1454.

Fernandez, A. A., & Bowser, P. R. (2010). Selection for a dominant oncogene and large male size as a risk factor for melanoma in the Xiphophorus animal model. Molecular Ecology, 19(15), 3114–3123.

Fernandez, A. A., & Morris, M. R. (2008). Mate choice for more melanin as a mechanism to maintain a functional oncogene. Proceedings of the National Academy of Sciences, 105(36), 13503–13507.

Formenti, G., Abueg, L., Brajuka, A., Brajuka, N., Gallardo-Alba, C., Giani, A., Fedrigo, O., & Jarvis, E. D. (2022). Gfastats: conversion, evaluation and manipulation of genome sequences using assembly graphs. Bioinformatics, 38(17), 4214–4216.

García-España, A., & Philips, M. R. (2023). Origin and evolution of RAS membrane targeting. Oncogene, 42(21), 1741–1750.

García-Navas, V., Kear, B. P., & Westerman, M. (2020). The geography of speciation in dasyurid marsupials. Journal of Biogeography, 47(9), 2042–2053.

Gemmell, R. T., Veitch, C., & Nelson, J. (2002). Birth in marsupials. Comparative Biochemistry and Physiology Part B: Biochemistry and Molecular Biology, 131(4), 621–630.

Goitre, L., Trapani, E., Trabalzini, L., & Retta, S. F. (2014). The Ras superfamily of small GTPases: the unlocked secrets. Ras signaling: methods and protocols, 1–18.

Guiler, E. (1970). Observations on the Tasmanian devil, Sarcophilus harrisii (Marsupialia: Dasyuridae) II. Reproduction, breeding and growth of pouch young. Australian Journal of Zoology, 18(1), 63–70.

Guindon, S., Dufayard, J.-F., Lefort, V., Anisimova, M., Hordijk, W., & Gascuel, O. (2010). New algorithms and methods to estimate maximum-likelihood phylogenies: assessing the performance of PhyML 3.0. Systematic Biology, 59(3), 307–321.

Hall, T. A. (1999). BioEdit: a user-friendly biological sequence alignment editor and analysis program for Windows 95/98/NT. Nucleic acids symposium series,

Hanahan, D. (2022). Hallmarks of cancer: new dimensions. Cancer discovery, 12(1), 31–46.

Harry, E. Paired REad TEXTure Viewer. OpenGL Powered Pretext Contact Map Viewer. In https://github.com/sanger-tol/PretextView

Hartley, G. A., Frankenberg, S. R., Robinson, N. M., MacDonald, A. J., Hamede, R. K., Burridge, C. P., Jones, M. E., Faulkner, T., Shute, H., & Rose, K. (2024). Genome of the endangered eastern quoll (Dasyurus viverrinus) reveals signatures of historical decline and pelage color evolution. Communications Biology, 7(1), 636.

Hartmann, K. (2012). Clinical aspects of feline retroviruses: a review. Viruses, 4(11), 2684–2710.

Harvey, J. (1964). An unidentified virus which causes the rapid production of tumours in mice. Nature, 204(4963), 1104–1105.

Hoang, D. T., Chernomor, O., Von Haeseler, A., Minh, B. Q., & Vinh, L. S. (2018). UFBoot2: improving the ultrafast bootstrap approximation. Molecular Biology and Evolution, 35(2), 518–522.

Hogg, C. J., Edwards, R. J., Farquharson, K. A., Silver, L. W., Brandies, P., Peel, E., Escalona, M., Jaya, F. R., Thavornkanlapachai, R., & Batley, K. (2024). Extant and extinct bilby genomes combined with Indigenous knowledge improve conservation of a unique Australian marsupial. Nature Ecology & Evolution, 8(7), 1311–1326.

Hopkins, D., & Gaynor, B. (1985). Schwannoma in a kowari (Dasyuroides byrnei). Australian veterinary journal, 62(10), 340–341. 10.1111/j.1751-0813.1985.tb07655.x

Ibeh, N., Feigin, C. Y., Frankenberg, S. R., McCarthy, D. J., Pask, A. J., & Romero, I. G. (2024). De novo transcriptome assembly and genome annotation of the fat-tailed dunnart (Sminthopsis crassicaudata). Gigabyte, 2024.

Jackson, S. (2007). Australian mammals: biology and captive management. Csiro Publishing.

Jeggo, P. A., Pearl, L. H., & Carr, A. M. (2016). DNA repair, genome stability and cancer: a historical perspective. Nature reviews cancer, 16(1), 35–42.

John, K., Alla, V., Meier, C., & Pützer, B. M. (2011). GRAMD4 mimics p53 and mediates the apoptotic function of p73 at mitochondria. Cell Death & Differentiation, 18(5), 874–886.

Kalyaanamoorthy, S., Minh, B. Q., Wong, T. K., Von Haeseler, A., & Jermiin, L. S. (2017). ModelFinder: fast model selection for accurate phylogenetic estimates. Nature methods, 14(6), 587–589.

Katoh, K., & Standley, D. M. (2013). MAFFT multiple sequence alignment software version 7: improvements in performance and usability. Molecular Biology and Evolution, 30(4), 772–780.

Keane, M., Semeiks, J., Webb, A. E., Li, Y. I., Quesada, V., Craig, T., Madsen, L. B., van Dam, S., Brawand, D., & Marques, P. I. (2015). Insights into the evolution of longevity from the bowhead whale genome. Cell reports, 10(1), 112–122.

Khalil, A., & Nemer, G. (2020). The potential oncogenic role of the RAS-like GTP-binding gene RIT1 in glioblastoma. Cancer Biomarkers, 29(4), 509–519.

Kirkwood, T. B. (1977). Evolution of ageing. Nature, 270(5635), 301–304.

Koh, J., Itahana, Y., Mendenhall, I. H., Low, D., Soh, E. X. Y., Guo, A. K., Chionh, Y. T., Wang, L.-F., & Itahana, K. (2019). ABCB1 protects bat cells from DNA damage induced by genotoxic compounds. Nature communications, 10(1), 1–14.

Kumar, S., Suleski, M., Craig, J. M., Kasprowicz, A. E., Sanderford, M., Li, M., Stecher, G., & Hedges, S. B. (2022). TimeTree 5: an expanded resource for species divergence times. Molecular Biology and Evolution, 39(8), msac174.

Ladds, P. (2009). 37. Neoplasia and Related Proliferations in Terrestrial Mammals. In Pathology of Australian native wildlife (pp. 429–457). BioOne.

Lariviere, D., Ostrovsky, A., Gallardo, C., Syme, A., Abueg, L., Pickett, B., Formenti, G., Sozzoni, M., & Nekrutenko, A. (2024). Vertebrate genome assembly using HiFi, Bionano and Hi-C data-Step by Step.

Letunic, I., Khedkar, S., & Bork, P. (2021). SMART: recent updates, new developments and status in 2020. Nucleic acids research, 49(D1), D458–D460.

Levitt, N. C., & Hickson, I. D. (2002). Caretaker tumour suppressor genes that defend genome integrity. Trends in molecular medicine, 8(4), 179–186.

Li, H. (2018). Minimap2: pairwise alignment for nucleotide sequences. Bioinformatics, 34(18), 3094–3100.

Liao, Y., Smyth, G. K., & Shi, W. (2014). featureCounts: an efficient general-purpose read summarization program. Bioinformatics, 30(7), 923–930.

Lima-de-Faria, A., Mitelman, F., Blomberg, J., & Pfeifer-Ohlsson, S. (1991). Telomeric location of retroviral oncogenes in humans. Hereditas, 114(3), 207–211.

Ljubuncic, P., & Reznick, A. Z. (2009). The evolutionary theories of aging revisited–a mini-review. Gerontology, 55(2), 205–216.

Lovell, J. T., Sreedasyam, A., Schranz, M. E., Wilson, M., Carlson, J. W., Harkess, A., Emms, D., Goodstein, D. M., & Schmutz, J. (2022). GENESPACE tracks regions of interest and gene copy number variation across multiple genomes. Elife, 11, e78526.

Lynch, M. (2008). 13. Bandicoots and bilbies. Medicine of Australian mammals, 439.

MacRae, S. L., Zhang, Q., Lemetre, C., Seim, I., Calder, R. B., Hoeijmakers, J., Suh, Y., Gladyshev, V. N., Seluanov, A., & Gorbunova, V. (2015). Comparative analysis of genome maintenance genes in naked mole rat, mouse, and human. Aging cell, 14(2), 288–291.

Madsen, T., Arnal, A., Vittecoq, M., Bernex, F., Abadie, J., Labrut, S., Garcia, D., Faugère, D., Lemberger, K., & Beckmann, C. (2017). Cancer prevalence and etiology in wild and captive animals. In Ecology and evolution of cancer (pp. 11–46). Elsevier.

Martin, M. (2011). Cutadapt removes adapter sequences from high-throughput sequencing reads. EMBnet. journal, 17(1), 10–12.

McEwen, G. K., Alquezar-Planas, D. E., Dayaram, A., Gillett, A., Tarlinton, R., Mongan, N., Chappell, K. J., Henning, J., Tan, M., & Timms, P. (2021). Retroviral integrations contribute to elevated host cancer rates during germline invasion. Nature communications, 12(1), 1316.

Mendes, F. K., Vanderpool, D., Fulton, B., & Hahn, M. W. (2020). CAFE 5 models variation in evolutionary rates among gene families. Bioinformatics, 36(22-23), 5516–5518.

Metzger, M. J., & Goff, S. P. (2016). A sixth modality of infectious disease: contagious cancer from devils to clams and beyond. PLoS Pathogens, 12(10), e1005904.

Minh, B. Q., Schmidt, H. A., Chernomor, O., Schrempf, D., Woodhams, M. D., Von Haeseler, A., & Lanfear, R. (2020). IQ-TREE 2: new models and efficient methods for phylogenetic inference in the genomic era. Molecular Biology and Evolution, 37(5), 1530–1534.

Morton, S., & Fletcher, T. (1989). Fauna of Australia.

Nunney, L., Maley, C. C., Breen, M., Hochberg, M. E., & Schiffman, J. D. (2015). Peto’s paradox and the promise of comparative oncology. In (Vol. 370, pp. 20140177): The Royal Society.

Obendorf, D. L. (1993). Diseases of dasyurid marsupials. The biology and management of Australasian carnivorous marsupials (M. Roberts, J. Carnio, G. Crawshaw, and M. Hutchins, eds.). Metropolitan Toronto Zoo and the Monotreme and Marsupial Advisory Group of the American Association of Zoological Parks and Aquariums, Toronto, Ontario, Canada, 39–46.

Old, J. M., & Deane, E. M. (2000). Development of the immune system and immunological protection in marsupial pouch young. Developmental & Comparative Immunology, 24(5), 445–454. 10.1016/S0145-305X(00)00008-2

Old, J. M., & Stannard, H. J. (2020). Conservation of quolls (Dasyurus spp.) in captivity–a review. Australian Mammalogy, 43(3), 277–289.

Pan, D., & Zhang, L. (2008). Tandemly arrayed genes in vertebrate genomes. International Journal of Genomics, 2008(1), 545269.

Parrott, M. L., & Edwards, A. M. (2023). Reproductive strategies and biology of the Australasian marsupials. In American and Australasian Marsupials: An Evolutionary, Biogeographical, and Ecological Approach (pp. 1–49). Springer.

Peck, S., Michael, S., Knowles, G., Davis, A., & Pemberton, D. (2019). Causes of mortality and severe morbidity requiring euthanasia in captive Tasmanian devils (Sarcophilus harrisii) in Tasmania. Australian veterinary journal, 97(4), 89–92.

Peel, E., Silver, L., Brandies, P., Zhu, Y., Cheng, Y., Hogg, C. J., & Belov, K. (2022). Best genome sequencing strategies for annotation of complex immune gene families in wildlife. GigaScience, 11, giac100.

Postler, T. S., Wang, A., Brundu, F. G., Wang, P., Wu, Z., Butler, K. E., Grinberg-Bleyer, Y., Krishnareddy, S., Lagana, S. M., & Saqi, A. (2023). A pan-cancer analysis implicates human NKIRAS1 as a tumor-suppressor gene. Proceedings of the National Academy of Sciences, 120(46), e2312595120.

Price, G., & Farquharson, K. . (2022). PacBio HiFi genome assembly using hifiasm v2.1. WorkflowHub. .

Prior, I. A., Lewis, P. D., & Mattos, C. (2012). A comprehensive survey of Ras mutations in cancer. Cancer research, 72(10), 2457–2467.

Quinlan, A. R., & Hall, I. M. (2010). BEDTools: a flexible suite of utilities for comparing genomic features. Bioinformatics, 26(6), 841–842.

Quinlan, M. P., & Settleman, J. (2009). Isoform-specific ras functions in development and cancer. Future oncology, 5(1), 105–116.

Rangwala, S. H., Kuznetsov, A., Ananiev, V., Asztalos, A., Borodin, E., Evgeniev, V., Joukov, V., Lotov, V., Pannu, R., & Rudnev, D. (2021). Accessing NCBI data using the NCBI sequence viewer and genome data viewer (GDV). Genome research, 31(1), 159–169.

Ratcliffe, H. L. (1933). Incidence and nature of tumors in captive wild mammals and birds. The American Journal of Cancer, 17(1), 116–135.

Renfree, M. B. (2010). Marsupials: placental mammals with a difference. Placenta, 31, S21–S26.

Rhie, A., McCarthy, S. A., Fedrigo, O., Damas, J., Formenti, G., Koren, S., Uliano-Silva, M., Chow, W., Fungtammasan, A., & Kim, J. (2021). Towards complete and error-free genome assemblies of all vertebrate species. Nature, 592(7856), 737–746.

Robinson, J. T., Thorvaldsdóttir, H., Winckler, W., Guttman, M., Lander, E. S., Getz, G., & Mesirov, J. P. (2011). Integrative genomics viewer. Nature biotechnology, 29(1), 24–26.

Roperto, S., Russo, V., Urraro, C., Restucci, B., Corrado, F., De Falco, F., & Roperto, F. (2017). ERas is constitutively expressed in full term placenta of pregnant cows. Theriogenology, 103, 162–168.

Ruprecht, K., Mayer, J., Sauter, M., Roemer, K., & Mueller-Lantzsch, N. (2008). Endogenous retroviruses: endogenous retroviruses and cancer. Cellular and Molecular Life Sciences, 65, 3366–3382.

Sanderson, M. J. (2003). r8s: inferring absolute rates of molecular evolution and divergence times in the absence of a molecular clock. Bioinformatics, 19(2), 301–302.

Santra, M. K., Wajapeyee, N., & Green, M. R. (2009). F-box protein FBXO31 mediates cyclin D1 degradation to induce G1 arrest after DNA damage. Nature, 459(7247), 722–725.

Schneider, V. A., Graves-Lindsay, T., Howe, K., Bouk, N., Chen, H.-C., Kitts, P. A., Murphy, T. D., Pruitt, K. D., Thibaud-Nissen, F., & Albracht, D. (2017). Evaluation of GRCh38 and de novo haploid genome assemblies demonstrates the enduring quality of the reference assembly. Genome research, 27(5), 849–864.

Simão, F. A., Waterhouse, R. M., Ioannidis, P., Kriventseva, E. V., & Zdobnov, E. M. (2015). BUSCO: assessing genome assembly and annotation completeness with single-copy orthologs. Bioinformatics, 31(19), 3210–3212.

Simmons, G., Young, P., Hanger, J., Jones, K., Clarke, D., McKee, J., & Meers, J. (2012). Prevalence of koala retrovirus in geographically diverse populations in Australia. Australian veterinary journal, 90(10), 404–409. 10.1111/j.1751-0813.2012.00964.x

Solovyev, V., Kosarev, P., Seledsov, I., & Vorobyev, D. (2006). Automatic annotation of eukaryotic genes, pseudogenes and promoters. Genome biology, 7, 1–12.

Sondka, Z., Dhir, N. B., Carvalho-Silva, D., Jupe, S., Madhumita, n., McLaren, K., Starkey, M., Ward, S., Wilding, J., & Ahmed, M. (2024). COSMIC: a curated database of somatic variants and clinical data for cancer. Nucleic acids research, 52(D1), D1210–D1217.

Spandidos, D. A., & Anderson, M. L. (1989). Oncogenes and onco-suppressor genes: their involvement in cancer. The Journal of pathology, 157(1), 1–10.

Stammnitz, M. R., Gori, K., Kwon, Y. M., Harry, E., Martin, F. J., Billis, K., Cheng, Y., Baez-Ortega, A., Chow, W., & Comte, S. (2023). The evolution of two transmissible cancers in Tasmanian devils. Science, 380(6642), 283–293.

Straube, E. F., & Callinan, R. B. (1980). Cutaneous squamous cell carcinoma associated with mammary adenocarcinoma in an eastern quoll Dasyurus viverrinus. Journal of Comparative Pathology, 90(3), 495–497. 10.1016/0021-9975(80)90020-1

Suarez-Cabrera, C., Ojeda-Perez, I., Sanchez-Baltasar, R., Page, A., Bravo, A., Navarro, M., & Ramirez, A. (2021). ERAS, a Member of the Ras Superfamily, Acts as an Oncoprotein in the Mammary Gland. Cancers, 13(21), 5588.

Sulak, M., Fong, L., Mika, K., Chigurupati, S., Yon, L., Mongan, N. P., Emes, R. D., & Lynch, V. J. (2016). TP53 copy number expansion is associated with the evolution of increased body size and an enhanced DNA damage response in elephants. Elife, 5, e11994.

Takahashi, K., Mitsui, K., & Yamanaka, S. (2003). Role of ERas in promoting tumour-like properties in mouse embryonic stem cells. Nature, 423(6939), 541–545.

Tanaka, Y., Ikeda, T., Kishi, Y., Masuda, S., Shibata, H., Takeuchi, K., Komura, M., Iwanaka, T., Muramatsu, S.-i., & Kondo, Y. (2009). ERas is expressed in primate embryonic stem cells but not related to tumorigenesis. Cell Transplantation, 18(4), 381–389.

Tarlinton, R., & Greenwood, A. D. (2024). Koala retrovirus and neoplasia: correlation and underlying mechanisms. Current Opinion in Virology, 67, 101427.

Tarlinton, R., Meers, J., Hanger, J., & Young, P. (2005). Real-time reverse transcriptase PCR for the endogenous koala retrovirus reveals an association between plasma viral load and neoplastic disease in koalas. Journal of General Virology, 86(3), 783–787.

Thies, K. A., Cole, M. W., Schafer, R. E., Spehar, J. M., Richardson, D. S., Steck, S. A., Das, M., Lian, A. W., Ray, A., & Shakya, R. (2021). The small G-protein RalA promotes progression and metastasis of triple-negative breast cancer. Breast cancer research, 23(1), 65.

Tian, R., Han, K., Geng, Y., Yang, C., Shi, C., Thomas, P. B., Pearce, C., Moffatt, K., Ma, S., & Xu, S. (2022). A chromosome-level genome of Antechinus flavipes provides a reference for an Australian marsupial genus with male death after mating. Molecular Ecology Resources, 22(2), 740–754.

Tollis, M., Robbins, J., Webb, A. E., Kuderna, L. F., Caulin, A. F., Garcia, J. D., Bèrubè, M., Pourmand, N., Marques-Bonet, T., & O’Connell, M. J. (2019). Return to the sea, get huge, beat cancer: an analysis of cetacean genomes including an assembly for the humpback whale (Megaptera novaeangliae). Molecular Biology and Evolution, 36(8), 1746–1763.

Twin, J. E., & Pearse, A. M. (1986). A malignant mixed salivary tumour and a mammary carcinoma in a young wild eastern spotted native cat Dasyurus viverrinus (marsupialia). Journal of Comparative Pathology, 96(3), 301–306. 10.1016/0021-9975(86)90050-2

Tyndale-Biscoe, C. H., & Renfree, M. (1987). Reproductive physiology of marsupials. Cambridge University Press.

Vincze, O., Colchero, F., Lemaître, J.-F., Conde, D. A., Pavard, S., Bieuville, M., Urrutia, A. O., Ujvari, B., Boddy, A. M., & Maley, C. C. (2022). Cancer risk across mammals. Nature, 601(7892), 263–267.

Wang, X., Douglas, K. C., VandeBerg, J. L., Clark, A. G., & Samollow, P. B. (2014). Chromosome-wide profiling of X-chromosome inactivation and epigenetic states in fetal brain and placenta of the opossum, Monodelphis domestica. Genome research, 24(1), 70–83.

Westerman, M., Krajewski, C., Kear, B. P., Meehan, L., Meredith, R. W., Emerling, C. A., & Springer, M. S. (2016). Phylogenetic relationships of dasyuromorphian marsupials revisited. Zoological Journal of the Linnean Society, 176(3), 686–701.

Wittbrodt, J., Adam, D., Malitschek, B., Mäueler, W., Raulf, F., Telling, A., Robertson, S. M., & Schartl, M. (1989). Novel putative receptor tyrosine kinase encoded by the melanoma-inducing Tu locus in Xiphophorus. Nature, 341(6241), 415–421.

Woolley, P. (1971). Maintenance and breeding of laboratory colonies. International Zoo Yearbook, 11(1), 351–354.

Zhang, Y., & Li, G. (2020). A tumor suppressor DLC1: The functions and signal pathways. Journal of cellular physiology, 235(6), 4999–5007.

Zhou, C., McCarthy, S. A., & Durbin, R. (2023). YaHS: yet another Hi-C scaffolding tool. Bioinformatics, 39(1), btac808.

