## Supplementary material for "When Cells Rebel: a comparative genomics investigation into marsupial cancer susceptibility"

### *1. Genome Assemblies*

The kowari genome was 3.21 GB and consisted of 851 scaffolds and 2,402 contigs. The scaffold N50 was 620 MB and the contig N50 was 4 MB. Benchmarking Universal Single-Copy Orthologs (BUSCO) v5.8.0 (Simão et al., 2015) identified 95.6% mammalian genes, of which 94.5% were single copy, 1.2% were duplicated, 1.0% were fragmented and 3.4% were missing.

The bandicoot genome was 3.94 GB and consisted of 1,462 scaffolds and 4,595 contigs. The scaffold N50 was 722 MB and the contig N50 was 2 MB. BUSCO v 5.8.0 identified 89.5% mammalian genes, of which 87.1% were single copy and 2.5% were duplicated. 1.5% were fragmented and 8.9% were missing.

We could not identify the SRY gene in either the kowari or the bandicoot. In addition, genome coverage appeared to be largely equal across the entire genome for both species, suggesting both samples game from a homogametic individual (Figure S1). Contact maps for both species indicated 7 large scaffolds, corresponding with the 7 pairs of chromosomes for both orders (Figure S2).

### *2. Genome Annotations*

BUSCO v5.8.0 (Simão et al., 2015) scores were calculated on the Galaxy Australia webserver (https://usegalaxy.org.au/) using the mammalia_odb10 database.

Repetitive regions of the genomes were masked using a 256 GB RAM, 64 vCPU, 3TB Pawsey Supercomputing Centre Nimbus cloud machine for the nine published genomes and the Galaxy webserver for the bandicoot and kowari. First, RepeatModeler v2.0.1 (Flynn et al., 2020) was used to identify and create a database of repetitive regions, then RepeatMasker v4.0.6 (Smit et al., 2013-2015) was used to mask them with the -nolow parameter to avoid masking low complexity regions.

A global assembly of all available tissue transcriptomes for each species was generated to guide annotation. Bioproject information for the raw data used for each genome is included in Supplementary Table 1. Raw reads were quality checked using FastQC v0.11.8 (Andrews, 2010) then trimmed using Trimmomatic v 0.39 (Bolger et al., 2014) with the parameters ILLUMINACLIP:TruSeq3-PE.fa:2:30:10, SLIDINGWINDOW:4:5, LEADING:5, TRAILING:5, MINLEN:25. Reads were aligned to the repeat masked genome using hisat2 v2.1.0 (Kim et al., 2019) and StringTie v2.1.6 (Pertea et al., 2015) was used to merge the aligned reads into tissue-specific transcriptomes. The transcriptomes were then merged into a global transcriptome using Stringtie *merge* and filtered to only include those with FPKM > 0.1 and length > 30bp and then TransDecoder v2.0.1 (Haas et al., 2013) was used to predict coding regions in the global transcriptome with a minimum length of 20 amino acids.

FGENESH++v7.2.2 (Solovyev et al., 2006) was used for genome annotation using the longest open reading frame predicted from the global transcriptome, mammalian settings and optimised parameters supplied with the Tasmanian devil (*Sarcophilus harrisii*) gene finding matrix.

*3. CAFE model selection*

We tried a number of different parameter combinations to determine the best model for the data. First, we determined whether to use a base model or a discrete gamma model with k=2 (meaning gene families can belong to one of two different evolutionary rate categories) or k=3. Each model was run five times to check for convergence. Any models that did not converge were rejected and the gamma model with k=2 was retained.

We then tried both a global $\lambda$ model and a multi-$\lambda$ model (with $\lambda_1$ for Australian marsupials and $\lambda_2$ for American marsupials). Again, both models were run five times to check for convergence. To determine if the more constrained global λ model was a better fit than the multi-λ model, we performed a likelihood ratio test. We ran 100 simulations under the null hypothesis (i.e. the global λ model). For each simulation, we calculated the likelihood of the global λ model ($L_{global}$), the likelihood of the multi-λ model ($L_{multi}$) and then used these values to calculate the likelihood ratio: $2(lnL_{global} - lnL_{multi})$. We plotted a histogram of the likelihood ratio of all the simulations and used this to determine the probability of obtaining the value of our actual likelihood ratio under the null hypothesis (Figure S7). As $p < 0.05$, we rejected the null hypothesis and selected the multi-λ model.

*Tables and Figures*

**Table S1.** Details for genomes used in this study

| Species | Common Name | Genome assembly (NCBI accession number) | Transcriptome source (NCBI project number) | Genome Size (GB) | Contigs | Scaffolds | Scaffold N50 (MB) | Contig N50 (MB) | Complete BUSCOs |
|---|---|---|---|---|---|---|---|---|---|
| *Dromiciops gliroides* | Monito del monte | mDroGli1.pri (GCF_019393635.1) | PRJNA416414 | 3.3 | 277 | 17 | 670.8 | 38.2 | 96.7% [Single copy:93.7% Duplicated:3.0%] |
| *Monodelphis domestica* | Gray short-tailed opossum | mMonDom1.pri (GCF_027887165.1) | PRJNA200320 | 3.6 | 2,268 | 13 | 538.3 | 3.9 | 94.9% [Single copy:92.5%, Duplicated:2.4%] |
| *Macrotis lagotis* | Greater bilby | bilby.v1.9 (GCF_037893015.1) | PRJNA1049866 | 3.7 | 5,027 | 608 | 343.9 | 1.2 | 95.6% [Single copy:90.8%, Duplicated:4.8%] |
| *Phascolarctos cinereus* | Koala | phaCin_HiC (GCA_003287225.2) | PRJNA230900, PRJNA327021 | 3.2 | 1,913 | 1,245 | 428.2 | 11.4 | 97.2% [Single copy:96.1%, Duplicated:1.1%] |
| *Notamacropus eugenii* | Tammar wallaby | Tammar_Male_v7_Final_Haploid_with_Y-001 | PRJDB1934 | 3.4 | 56 | 9 | 483.6 | 194.0 | 96.9% [Single copy:94.7%, Duplicated:2.3%] |
| *Sminthopsis crassicaudata* | Fat-tailed dunnart | dunnart_asm_12-2021_sm (Figshare) | PRJNA1028148 | 3.2 | 2,569 | 1,848 | 72.6 | 11.2 | 96.4% [Single copy:94.9%, Duplicated:1.5%] |
| *Antechinus flavipes* | Yellow-footed antechinus | AdamAnt_v2 (GCF_016432865.1) | PRJNA565840 | 3.2 | 1,103 | 485 | 636.7 | 51.8 | 94.9% [Single copy:93.7%, Duplicated:1.2%] |
| *Sarcophilus harrisii* | Tasmanian devil | mSarHar1.11 (GCF_902635505.1) | PRJEB34650 | 3.1 | 444 | 105 | 611.3 | 62.3 | 94.7% [Single copy:93.6%, Duplicated:1.1%] |
| *Dasyurus viverrinus* | Eastern quoll | UniMelb_DasViv_v1.0 (GCA_020854095.1) | PRJNA963007 | 3.1 | 507 | 76 | 628.5 | 13.8 | 96.4% [Single copy:95.4%, Duplicated:1.0%] |
| *Dasyuroides byrneii* | Kowari | mDasByr.1.2_20251403 | this study | 3.2 | 2,402 | 851 | 620.0 | 4.0 | 95.6% [Single copy:94.5%, Duplicated:1.2%] |
| *Perameles gunnii* | Eastern barred bandicoot | mPerGun1.2_20251003 | this study | 3.9 | 4,595 | 1,462 | 722.0 | 2.0 | 89.5% [Single copy:87.1%, Duplicated:2.5%] |

**Table S2**. Reported cases of neoplasia across marsupial taxa. Species are grouped by family, with the exception of Peramelemorphia and Phalangeriformes, as species in these taxa are often grouped together in reporting. ** indicates that this figure may not include all captive devils.

| ***Taxon*** | ***Common name*** | ***Ratcliffe, 1933*** | ***Effron, Griner & Benirschke, 1977*** | ***Canfield & Cunningham, 1993*** | ***Canfield Hartley & Reddacliff, 1990a; Canfield, Hartley, & Reddacliff, 1990b*** | ***Ladds, 2009*** | ***Wildlife Health Registry (2004-2024)*** | ***Total*** |
|---|---|---|---|---|---|---|---|---|
| *Dasyuridae* | *Dasyurids* | *3* | *3* | *7* | *70* | *140* | *59*** | *282* |
| *Didelphidae* | *Opossums* | *3* | *1* | *not reported* | *not reported* | *not reported* | *not reported* | *4* |
| *Macropodidae* | *Kangaroos, wallabies, quokkas* | *2* | *4* | *11* | *14* | *43* | *17* | *91* |
| *Peramelemorphia* | *Bandicoots and bilbies* | *1* | *not reported* | *not reported* | *3* | *19* | *12* | *35* |
| *Phascolarctidae* | *Koala* | *not reported* | *not reported* | *not reported* | *26* | *70* | *113* | *209* |
| *Phalangeriformes* | *Possums and gliders* | *not reported* | *not reported* | *0* | *22* | *74* | *33* | *129* |
| *Potoroidae* | *Bettongs, potoroos* | *not reported* | *not reported* | *not reported* | *not reported* | *not reported* | *3* | *3* |
| *Vombatidae* | *Wombats* | *not reported* | *not reported* | *0* | *2* | *2* | *not reported* | *4* |

**Table S3**. Reported cases of neoplasia across dasyurid genera. ** indicates that this figure may not include all captive devils.

| Genus | Common name | Ratcliffe, 1933 | Effron, Griner & Benirschke, 1977 | Canfield, Hartley, & Reddacliff, 1990b | Ladds, 2009 | Wildlife Health Registry (2004-2024)** | Total |
|---|---|---|---|---|---|---|---|
| Antechinus | Antechinus | not reported | not reported | 4 | 10 | 1 | 15 |
| Dasycercus | Mulgara | not reported | not reported | 1 | 2 | 1 | 4 |
| Dasyuroides | Kowari | not reported | not reported | 11 | 32 | not reported | 43 |
| Dasyurus | Quoll | 2 | not reported | 20 | 45 | 12 | 79 |
| Parantechinus | Dibbler | not reported | not reported | 3 | not reported | not reported | 3 |
| Phascogale | Phascogale | not reported | not reported | 3 | 12 | 1 | 16 |
| Planigale | Planigale | not reported | not reported | 4 | 4 | not reported | 8 |
| Pseudoantechinus | False antechinus | not reported | not reported | 5 | not reported | not reported | 5 |
| Sarcophilus | Devil | 1 | 3 | 16 | 26 | 43** | 89 |
| Sminthopsis | Dunnart | not reported | not reported | 3 | 8 | 1 | 12 |

**Table S1.** Number of protein-coding genes annotated by FGENESH in the 11 species, number annotated as cancer orthologs (Cosmic database), and percentage assigned to orthogroups.

| Species | Total genes annotated by FGENESH | Annotated as cancer orthologs | Total genes remaining after filtering | Percentage of Genes in Orthogroups |
|---|---|---|---|---|
| Yellow footed antechinus | 36,780 | 646 | 25,727 | 86.3 |
| Eastern Barred Bandicoot | 76,963 | 627 | 38,979 | 84.5 |
| Greater Bilby | 37,266 | 622 | 28,951 | 81.2 |
| Tasmanian devil | 32,130 | 648 | 24,172 | 83.4 |
| Fat tailed dunnart | 33,770 | 662 | 24,285 | 85.3 |
| Koala | 28,365 | 672 | 23,515 | 86.2 |
| Kowari | 73,135 | 674 | 39,773 | 88.9 |
| Monito del Monte | 28,719 | 588 | 23,917 | 85.7 |
| Gray Short Tailed Opossum | 36,640 | 576 | 30,251 | 68.8 |
| Eastern Quoll | 32,702 | 640 | 22,463 | 91.2 |
| Tammar Wallaby | 34,784 | 649 | 26,687 | 87.4 |

**Table S2**. Divergence dates for nodes in marsupial phylogeny that were obtained from timetree.org

| Node | Time (million years ago) |
|---|---|
| Quoll, devil | 9.4 |
| Kowari, (quoll and devil) | 13.9 |
| Antechinus, dasyurini | 18 |
| Dunnart, (antechinus, kowari, quoll, devil) | 24.6 |
| Bilby, bandicoot | 30 |
| Bilby, dasyurids | 58 |
| Koala, tammar | 53 |
| Bilby, diprotodontia | 61 |
| Monito, eomarsupialia | 63 |
| Monito, opossum | 78 |

**Table S3.** Query sequences used for RAS annotations

| Species | Gene | Accession (Uniprot) |
|---|---|---|
| Homo sapiens | KRAS4A | P01116 |
| Homo sapiens | KRAS4B | P01116-2 |
| Homo sapiens | NRAS | P01111 |
| Homo sapiens | HRAS | P01112 |
| Homo sapiens | ERAS | Q7Z444 |
| Homo sapiens | RRAS | P10301 |
| Homo sapiens | RRAS2 | P62070 |
| Homo sapiens | MRAS | O14807 |
| Homo sapiens | RIT1 | Q92963 |
| Homo sapiens | RIT2 | Q99578 |
| Homo sapiens | RAP1A | P62834 |
| Homo sapiens | RAP1B | P61224 |
| Homo sapiens | RAP2A | P10114 |
| Homo sapiens | RAP2C | Q9Y3L5 |
| Homo sapiens | RAP2B | P61225 |
| Homo sapiens | RALA | P11233 |
| Homo sapiens | RALB | P11234 |
| Homo sapiens | REM1 | O75628 |
| Homo sapiens | REM2 | Q8IYK8 |
| Homo sapiens | RRAD | P55042 |
| Homo sapiens | GEM | P55040 |
| Homo sapiens | RERG | Q96A58 |
| Homo sapiens | RASL11A | Q6T310 |
| Homo sapiens | RASL11B | Q9BPW5 |
| Homo sapiens | DIRAS1 | O95057 |
| Homo sapiens | RASL10A | Q92737 |
| Homo sapiens | NKIRAS1 | Q9NYS0 |
| Homo sapiens | RASL12 | Q9NYN1 |
| Homo sapiens | RERGL | Q9H628 |
| Homo sapiens | RHEB | Q15382 |
| Homo sapiens | RHEBL1 | Q8TAI7 |
| Homo sapiens | DIRAS3 | O95661 |
| Homo sapiens | DIRAS2 | Q96HU8 |
| Homo sapiens | RASL10B | Q96S79 |
| Homo sapiens | NKIRAS2 | Q9NYR9 |
| Homo sapiens | RASD2 | Q96D21 |
| Homo sapiens | RASD1 | Q9Y272 |

**Table S4.** Accession numbers and gene names for nodes in the phylogenetic tree

| Accession number | Gene | Species | Phylo_Group | Gene_Subfamily |
|---|---|---|---|---|
| NM_181548.2 | >ERAS_MOUSE | House mouse | Eutherian | ERAS |
| XM_059883764.1 | >ERAS_COW | Domestic cattle | Eutherian | ERAS |
| XM_038448950.1 | >ERAS_DOG | Dog | Eutherian | ERAS |
| XM_005878010.2 | >ERAS_BAT | Brandt's bat | Eutherian | ERAS |
| XM_008272640.2 | >ERAS_RABBIT | Rabbit | Eutherian | ERAS |
| XM_004690133.1 | >ERAS_MOLE | Star-nosed mole | Eutherian | ERAS |
| XM_070258069.1 | >ERAS_HORSE | Horse | Eutherian | ERAS |
| XM_036917191.2 | >ERAS_PANGOLIN | Chinese pangolin | Eutherian | ERAS |
| XM_008571436.1 | >ERAS_FLYING_LEMUR | Sunda flying lemur | Eutherian | ERAS |
| XM_006171940.1 | >ERAS_TREE_SHREW | Chinese tree shrew | Eutherian | ERAS |
| XM_003417997.3 | >ERAS_ELEPHANT | African savanna elephant | Eutherian | ERAS |
| XM_004464982.2 | >ERAS_ARMADILLO | Nine-banded armadillo | Eutherian | ERAS |
| XM_004464982.2 | >ERAS_SLOTH | Southern two-toed sloth | Eutherian | ERAS |
| NM_181532.3 | >ERAS_HUMAN | Homo sapiens | Eutherian | ERAS |
| NG_042222.1 | >RRAS_HUMAN | Homo sapiens | Eutherian | Ras subfamily |
| NG_017058.1 | >RRAS2_HUMAN | Homo sapiens | Eutherian | Ras subfamily |
| NM_001085049.3 | >MRAS_HUMAN | Homo sapiens | Eutherian | Ras subfamily |
| NG_033885.1 | >RIT1_HUMAN | Homo sapiens | Eutherian | Ras subfamily |
| NM_001272077.2 | >RIT2_HUMAN | Homo sapiens | Eutherian | Ras subfamily |
| NM_001010935.3 | >RAP1A_HUMAN | Homo sapiens | Eutherian | Ras subfamily |
| NM_001010942.3 | >RAP1B_HUMAN | Homo sapiens | Eutherian | Ras subfamily |
| NM_021033.7 | >RAP2A_HUMAN | Homo sapiens | Eutherian | Ras subfamily |
| NM_001271186.2 | >RAP2C_HUMAN | Homo sapiens | Eutherian | Ras subfamily |
| NM_002886.4 | >RAP2B_HUMAN | Homo sapiens | Eutherian | Ras subfamily |
| NM_005402.4 | >RALA_HUMAN | Homo sapiens | Eutherian | Ras subfamily |
| NM_001369400.1 | >RALB_HUMAN | Homo sapiens | Eutherian | Ras subfamily |
| NG_046939.1 | >REM1_HUMAN | Homo sapiens | Eutherian | Ras subfamily |
| NM_173527.3 | >REM2_HUMAN | Homo sapiens | Eutherian | Ras subfamily |
| NM_001128850.2 | >RRAD_HUMAN | Homo sapiens | Eutherian | Ras subfamily |
| NM_005261.4 | >GEM_HUMAN | Homo sapiens | Eutherian | Ras subfamily |
| NM_001190726.2 | >RERG_HUMAN | Homo sapiens | Eutherian | Ras subfamily |
| NM_001331126.2 | >RASL11A_HUMAN | Homo sapiens | Eutherian | Ras subfamily |
| NM_023940.3 | >RASL11b_HUMAN | Homo sapiens | Eutherian | Ras subfamily |
| NM_145173.4 | >DIRAS1_HUMAN | Homo sapiens | Eutherian | Ras subfamily |
| NM_006477.5 | >RASL10A_HUMAN | Homo sapiens | Eutherian | Ras subfamily |
| NM_001377351.1 | >NKIRAS1_HUMAN | Homo sapiens | Eutherian | Ras subfamily |
| NM_001307930.2 | >RASL12_HUMAN | Homo sapiens | Eutherian | Ras subfamily |
| NG_052618.1 | >RERGL_HUMAN | Homo sapiens | Eutherian | Ras subfamily |
| NM_005614.4 | >RHEB_HUMAN | Homo sapiens | Eutherian | Ras subfamily |
| NM_001303126.2 | >RHEBL1_HUMAN | Homo sapiens | Eutherian | Ras subfamily |
| NG_011753.1 | >DIRAS3_HUMAN | Homo sapiens | Eutherian | Ras subfamily |
| NM_017594.5 | >DIRAS2_HUMAN | Homo sapiens | Eutherian | Ras subfamily |
| NM_033315.4 | >RASL10b_HUMAN | Homo sapiens | Eutherian | Ras subfamily |
| NM_001001349.2 | >NKIRAS2_HUMAN | Homo sapiens | Eutherian | Ras subfamily |
| NM_001366725.1 | >RASD2_HUMAN | Homo sapiens | Eutherian | Ras subfamily |
| NG_028074.2 | >RASD1_HUMAN | Homo sapiens | Eutherian | Ras subfamily |
| NM_001369786.1 | >KRAS4A_HUMAN | Homo sapiens | Eutherian | Canonical |
| NM_001369787.1 | >KRAS4B_HUMAN | Homo sapiens | Eutherian | Canonical |

| Accession number | Gene | Species | Phylo_Group | Gene_Subfamily |
|---|---|---|---|---|
| NG_007572.1 | >NRAS_HUMAN | Homo sapiens | Eutherian | Canonical |
| NG_007666.1 | >HRAS_HUMAN | Homo sapiens | Eutherian | Canonical |
| XM_064499648.1 | >RRAS_BIRD | Emu | Aves | Ras subfamily |
| NM_204489.2 | >MRAS_BIRD | Chicken | Aves | Ras subfamily |
| NM_001031327.3 | >RIT1_BIRD | Chicken | Aves | Ras subfamily |
| XM_001233996.7 | >RIT2_BIRD | Chicken | Aves | Ras subfamily |
| XM_046904296.1 | >RAP1A_BIRD | Chicken | Aves | Ras subfamily |
| NM_001007852.1 | >RAP1B_BIRD | Chicken | Aves | Ras subfamily |
| XM_001233103.6 | >RAP2A_BIRD | Chicken | Aves | Ras subfamily |
| NM_001012572.3 | >RAP2C_BIRD | Chicken | Aves | Ras subfamily |
| XM_015291825.4 | >RAP2B_BIRD | Chicken | Aves | Ras subfamily |
| XM_046937036.1 | >RALA_BIRD | Chicken | Aves | Ras subfamily |
| XM_025152428.3 | >RALB_BIRD | Chicken | Aves | Ras subfamily |
| XM_015296468.4 | >REM1_BIRD | Chicken | Aves | Ras subfamily |
| XM_064475528.1 | >REM2_BIRD | Great cormorant | Aves | Ras subfamily |
| NM_001277606.3 | >RRAD_BIRD | Chicken | Aves | Ras subfamily |
| NM_213579.2 | >GEM_BIRD | Chicken | Aves | Ras subfamily |
| XM_046904998.1 | >RERG_BIRD | Chicken | Aves | Ras subfamily |
| XM_417126.8 | >RASL11A_BIRD | Chicken | Aves | Ras subfamily |
| XM_420710.8 | >RASL11b_BIRD | Chicken | Aves | Ras subfamily |
| XM_015299968.4 | >DIRAS1_BIRD | Chicken | Aves | Ras subfamily |
| NM_001030702.2 | >RASL10A_BIRD | Chicken | Aves | Ras subfamily |
| XM_040696256.2 | >NKIRAS1_BIRD | Chicken | Aves | Ras subfamily |
| XM_004943886.5 | >RASL12_BIRD | Chicken | Aves | Ras subfamily |
| XM_416411.8 | >RERGL_BIRD | Chicken | Aves | Ras subfamily |
| XM_040695391.2 | >RHEB_BIRD | Chicken | Aves | Ras subfamily |
| XM_065043780.1 | >RHEBL1_BIRD | Rock pigeon | Aves | Ras subfamily |
| XM_021296473.2 | >DIRAS3_BIRD | Rock pigeon | Aves | Ras subfamily |
| XM_423026.8 | >DIRAS2_BIRD | Chicken | Aves | Ras subfamily |
| XM_001233673.7 | >RASL10b_BIRD | Chicken | Aves | Ras subfamily |
| NM_001006333.2 | >NKIRAS2_BIRD | Chicken | Aves | Ras subfamily |
| XM_416293.8 | >RASD2_BIRD | Chicken | Aves | Ras subfamily |
| NM_001044636.2 | >RASD1_BIRD | Chicken | Aves | Ras subfamily |
| NM_001256162.1 | >KRAS_BIRD | Chicken | Aves | Canonical |
| NM_001012549.2 | >NRAS_BIRD | Chicken | Aves | Canonical |
| NM_001396746.1 | >HRAS_BIRD | Chicken | Aves | Canonical |
| XM_033171113.1 | >KRASBL_LIZARD | Sand lizard | Lizard | Canonical |
| NM_001114248.1 | >RRAS_FROG | Tropical clawed frog | Amphibian | Ras subfamily |
| XM_031900379.1 | >RRAS2_FROG | Tropical clawed frog | Amphibian | Ras subfamily |
| XM_002943157.5 | >MRAS_FROG | Tropical clawed frog | Amphibian | Ras subfamily |
| XM_031891490.1 | >RIT1_FROG | Tropical clawed frog | Amphibian | Ras subfamily |
| NM_001102887.1 | >RAP1A_FROG | Tropical clawed frog | Amphibian | Ras subfamily |
| NM_001008194.2 | >RAP1B_FROG | Tropical clawed frog | Amphibian | Ras subfamily |
| NM_001035116.1 | >RAP2A_FROG | Tropical clawed frog | Amphibian | Ras subfamily |
| NM_001016898.2 | >RAP2C_FROG | Tropical clawed frog | Amphibian | Ras subfamily |
| NM_001035115.1 | >RAP2B_FROG | Tropical clawed frog | Amphibian | Ras subfamily |
| NM_001015915.2 | >RALA_FROG | Tropical clawed frog | Amphibian | Ras subfamily |
| NM_001102845.1 | >RALB_FROG | Tropical clawed frog | Amphibian | Ras subfamily |
| NM_001102791.1 | >REM1_FROG | Tropical clawed frog | Amphibian | Ras subfamily |

| Accession number | Gene | Species | Phylo_Group | Gene_Subfamily |
|---|---|---|---|---|
| XM_002941509.5 | >REM2_FROG | Tropical clawed frog | Amphibian | Ras subfamily |
| NM_001016726.2 | >RRAD_FROG | Tropical clawed frog | Amphibian | Ras subfamily |
| NM_001097353.1 | >GEM_FROG | Tropical clawed frog | Amphibian | Ras subfamily |
| XM_040345529.1 | >RERG_FROG | Common frog | Amphibian | Ras subfamily |
| XM_002941535.4 | >RASL11A_FROG | Tropical clawed frog | Amphibian | Ras subfamily |
| NM_001015774.1 | >RASL11b_FROG | Tropical clawed frog | Amphibian | Ras subfamily |
| NM_001079312.1 | >DIRAS1_FROG | Tropical clawed frog | Amphibian | Ras subfamily |
| NM_001005037.1 | >RASL10A_FROG | Tropical clawed frog | Amphibian | Ras subfamily |
| XM_002932411.5 | >NKIRAS1_FROG | Tropical clawed frog | Amphibian | Ras subfamily |
| XM_004912687.4 | >RASL12_FROG | Tropical clawed frog | Amphibian | Ras subfamily |
| XM_040345486.1 | >RERGL_FROG | Common frog | Amphibian | Ras subfamily |
| NM_001015922.2 | >RHEB_FROG | Tropical clawed frog | Amphibian | Ras subfamily |
| NM_203606.2 | >RHEBL1_FROG | Tropical clawed frog | Amphibian | Ras subfamily |
| XM_031899814.1 | >DIRAS3_FROG | Tropical clawed frog | Amphibian | Ras subfamily |
| XM_002937026.4 | >DIRAS2_FROG | Tropical clawed frog | Amphibian | Ras subfamily |
| XM_004911629.4 | >RASL10b_FROG | Tropical clawed frog | Amphibian | Ras subfamily |
| XM_002939033.5 | >NKIRAS2_FROG | Tropical clawed frog | Amphibian | Ras subfamily |
| NM_001016006.2 | >RASD2_FROG | Tropical clawed frog | Amphibian | Ras subfamily |
| NM_001078940.1 | >RASD1_FROG | Tropical clawed frog | Amphibian | Ras subfamily |
| XM_040437469.1 | >KRAS_TOAD | Common toad | Amphibian | Canonical |
| NM_001016763.2 | >NRAS_FROG | Tropical clawed frog | Amphibian | Canonical |
| NM_001017003.2 | >HRAS_FROG | Tropical clawed frog | Amphibian | Canonical |
| NM_001008033.1 | >KRASBL_FROG | Tropical clawed frog | Amphibian | Canonical |
| NM_001005931.2 | >RRAS_FISH | Zebrafish | Fish | Ras subfamily |
| NM_001017815.1 | >RRAS2_FISH | Zebrafish | Fish | Ras subfamily |
| XM_033024843.1 | >MRAS_FISH | Thorny skate | Fish | Ras subfamily |
| NM_001128781.1 | >RIT1_FISH | Thorny skate | Fish | Ras subfamily |
| XM_033031348.1 | >RIT2_FISH | Thorny skate | Fish | Ras subfamily |
| NM_001002152.1 | >RAP1A_FISH | Zebrafish | Fish | Ras subfamily |
| NM_199533.1 | >RAP1B_FISH | Zebrafish | Fish | Ras subfamily |
| NM_001145705.1 | >RAP2A_FISH | Zebrafish | Fish | Ras subfamily |
| NM_001007055.1 | >RAP2C_FISH | Zebrafish | Fish | Ras subfamily |
| NM_001001729.2 | >RAP2B_FISH | Zebrafish | Fish | Ras subfamily |
| NM_201018.1 | >RALA_FISH | Zebrafish | Fish | Ras subfamily |
| NM_001003649.1 | >RALB_FISH | Zebrafish | Fish | Ras subfamily |
| NM_201174.1 | >REM1_FISH | Zebrafish | Fish | Ras subfamily |
| NM_001123046.1 | >REM2_FISH | Zebrafish | Fish | Ras subfamily |
| NM_199798.1 | >RRAD_FISH | Zebrafish | Fish | Ras subfamily |
| NM_001045849.1 | >GEM_FISH | Zebrafish | Fish | Ras subfamily |
| NM_001327837.1 | >RERG_FISH | Zebrafish | Fish | Ras subfamily |
| NM_001017840.2 | >RASL11A_FISH | Zebrafish | Fish | Ras subfamily |
| NM_200140.1 | >RASL11b_FISH | Zebrafish | Fish | Ras subfamily |
| NM_199831.1 | >DIRAS1_FISH | Zebrafish | Fish | Ras subfamily |
| NM_001128366.1 | >RASL10A_FISH | Zebrafish | Fish | Ras subfamily |
| NM_001100076.1 | >NKIRAS1_FISH | Zebrafish | Fish | Ras subfamily |
| NM_200395.1 | >RASL12_FISH | Zebrafish | Fish | Ras subfamily |
| NM_001002494.1 | >RERGL_FISH | Zebrafish | Fish | Ras subfamily |
| NM_200729.1 | >RHEB_FISH | Zebrafish | Fish | Ras subfamily |
| NM_001076748.2 | >RHEBL1_FISH | Zebrafish | Fish | Ras subfamily |

| Accession number | Gene | Species | Phylo_Group | Gene_Subfamily |
|---|---|---|---|---|
| XM_033028462.1 | >DIRAS3_FISH | Thorny skate | Fish | Ras subfamily |
| XM_005155552.4 | >DIRAS2_FISH | Zebrafish | Fish | Ras subfamily |
| XM_068221542.1 | >RASL10b_FISH | Zebrafish | Fish | Ras subfamily |
| NM_001003433.1 | >NKIRAS2_FISH | Zebrafish | Fish | Ras subfamily |
| NM_001030202.2 | >RASD2_FISH | Zebrafish | Fish | Ras subfamily |
| NM_200532.1 | >RASD1_FISH | Zebrafish | Fish | Ras subfamily |
| NM_001003744.2 | >KRAS_FISH | Zebrafish | Fish | Canonical |
| NM_131145.2 | >NRAS_FISH | Zebrafish | Fish | Canonical |
| NM_001017623.1 | >HRAS_FISH | Zebrafish | Fish | Canonical |
| NM_001292570.1 | >KRASBL_SHARK | Elephant Shark | Fish | Canonical |
| This study | >Af_HRas | Yellow-footed antechinus | Dasyuromorphia | Canonical |
| This study | >Af_KRas4a | Yellow-footed antechinus | Dasyuromorphia | Canonical |
| This study | >Af_KRas4b | Yellow-footed antechinus | Dasyuromorphia | Canonical |
| This study | >Af_NRas | Yellow-footed antechinus | Dasyuromorphia | Canonical |
| This study | >AfMgRas1 | Yellow-footed antechinus | Dasyuromorphia | MgRas |
| This study | >AfMgRas10 | Yellow-footed antechinus | Dasyuromorphia | MgRas |
| This study | >AfMgRas11 | Yellow-footed antechinus | Dasyuromorphia | MgRas |
| This study | >AfMgRas12 | Yellow-footed antechinus | Dasyuromorphia | MgRas |
| This study | >AfMgRas13 | Yellow-footed antechinus | Dasyuromorphia | MgRas |
| This study | >AfMgRas14 | Yellow-footed antechinus | Dasyuromorphia | MgRas |
| This study | >AfMgRas15 | Yellow-footed antechinus | Dasyuromorphia | MgRas |
| This study | >AfMgRas16 | Yellow-footed antechinus | Dasyuromorphia | MgRas |
| This study | >AfMgRas17 | Yellow-footed antechinus | Dasyuromorphia | MgRas |
| This study | >AfMgRas18 | Yellow-footed antechinus | Dasyuromorphia | MgRas |
| This study | >AfMgRas19 | Yellow-footed antechinus | Dasyuromorphia | MgRas |
| This study | >AfMgRas2 | Yellow-footed antechinus | Dasyuromorphia | MgRas |
| This study | >AfMgRas20 | Yellow-footed antechinus | Dasyuromorphia | MgRas |
| This study | >AfMgRas21 | Yellow-footed antechinus | Dasyuromorphia | MgRas |
| This study | >AfMgRas22 | Yellow-footed antechinus | Dasyuromorphia | MgRas |
| This study | >AfMgRas23 | Yellow-footed antechinus | Dasyuromorphia | MgRas |
| This study | >AfMgRas3 | Yellow-footed antechinus | Dasyuromorphia | MgRas |
| This study | >AfMgRas4 | Yellow-footed antechinus | Dasyuromorphia | MgRas |
| This study | >AfMgRas5 | Yellow-footed antechinus | Dasyuromorphia | MgRas |
| This study | >AfMgRas6 | Yellow-footed antechinus | Dasyuromorphia | MgRas |
| This study | >AfMgRas7 | Yellow-footed antechinus | Dasyuromorphia | MgRas |
| This study | >AfMgRas8 | Yellow-footed antechinus | Dasyuromorphia | MgRas |
| This study | >AfMgRas9 | Yellow-footed antechinus | Dasyuromorphia | MgRas |
| This study | >Db_Hras | Kowari | Dasyuromorphia | Canonical |
| This study | >Db_Kras4A | Kowari | Dasyuromorphia | Canonical |
| This study | >Db_Kras4B | Kowari | Dasyuromorphia | Canonical |
| This study | >Db_Nras | Kowari | Dasyuromorphia | Canonical |
| This study | >DbMgRas1 | Kowari | Dasyuromorphia | MgRas |
| This study | >DbMgRas10 | Kowari | Dasyuromorphia | MgRas |
| This study | >DbMgRas11 | Kowari | Dasyuromorphia | MgRas |
| This study | >DbMgRas12 | Kowari | Dasyuromorphia | MgRas |
| This study | >DbMgRas13 | Kowari | Dasyuromorphia | MgRas |
| This study | >DbMgRas14 | Kowari | Dasyuromorphia | MgRas |
| This study | >DbMgRas15 | Kowari | Dasyuromorphia | MgRas |
| This study | >DbMgRas16 | Kowari | Dasyuromorphia | MgRas |

| Accession number | Gene | Species | Phylo_Group | Gene_Subfamily |
|---|---|---|---|---|
| This study | >DbMgRas17 | Kowari | Dasyuromorphia | MgRas |
| This study | >DbMgRas18 | Kowari | Dasyuromorphia | MgRas |
| This study | >DbMgRas19 | Kowari | Dasyuromorphia | MgRas |
| This study | >DbMgRas2 | Kowari | Dasyuromorphia | MgRas |
| This study | >DbMgRas20 | Kowari | Dasyuromorphia | MgRas |
| This study | >DbMgRas21 | Kowari | Dasyuromorphia | MgRas |
| This study | >DbMgRas3 | Kowari | Dasyuromorphia | MgRas |
| This study | >DbMgRas4 | Kowari | Dasyuromorphia | MgRas |
| This study | >DbMgRas5 | Kowari | Dasyuromorphia | MgRas |
| This study | >DbMgRas6 | Kowari | Dasyuromorphia | MgRas |
| This study | >DbMgRas7 | Kowari | Dasyuromorphia | MgRas |
| This study | >DbMgRas8 | Kowari | Dasyuromorphia | MgRas |
| This study | >DbMgRas9 | Kowari | Dasyuromorphia | MgRas |
| This study | >Dg_HRas | Monito del monte | Microbiotheria | Canonical |
| This study | >Dg_KRas4A | Monito del monte | Microbiotheria | Canonical |
| This study | >Dg_KRas4B | Monito del monte | Microbiotheria | Canonical |
| This study | >Dg_NRas | Monito del monte | Microbiotheria | Canonical |
| This study | >DgMgRas1 | Monito del monte | Microbiotheria | MgRas |
| This study | >DgMgRas2 | Monito del monte | Microbiotheria | MgRas |
| This study | >Dv_HRas | Eastern quoll | Dasyuromorphia | Canonical |
| This study | >Dv_KRas4A | Eastern quoll | Dasyuromorphia | Canonical |
| This study | >Dv_KRas4B | Eastern quoll | Dasyuromorphia | Canonical |
| This study | >Dv_NRas | Eastern quoll | Dasyuromorphia | Canonical |
| This study | >DvMgRas1 | Eastern quoll | Dasyuromorphia | MgRas |
| This study | >DvMgRas10 | Eastern quoll | Dasyuromorphia | MgRas |
| This study | >DvMgRas11 | Eastern quoll | Dasyuromorphia | MgRas |
| This study | >DvMgRas12 | Eastern quoll | Dasyuromorphia | MgRas |
| This study | >DvMgRas13 | Eastern quoll | Dasyuromorphia | MgRas |
| This study | >DvMgRas14 | Eastern quoll | Dasyuromorphia | MgRas |
| This study | >DvMgRas15 | Eastern quoll | Dasyuromorphia | MgRas |
| This study | >DvMgRas16 | Eastern quoll | Dasyuromorphia | MgRas |
| This study | >DvMgRas17 | Eastern quoll | Dasyuromorphia | MgRas |
| This study | >DvMgRas18 | Eastern quoll | Dasyuromorphia | MgRas |
| This study | >DvMgRas19 | Eastern quoll | Dasyuromorphia | MgRas |
| This study | >DvMgRas2 | Eastern quoll | Dasyuromorphia | MgRas |
| This study | >DvMgRas20 | Eastern quoll | Dasyuromorphia | MgRas |
| This study | >DvMgRas21 | Eastern quoll | Dasyuromorphia | MgRas |
| This study | >DvMgRas22 | Eastern quoll | Dasyuromorphia | MgRas |
| This study | >DvMgRas3 | Eastern quoll | Dasyuromorphia | MgRas |
| This study | >DvMgRas4 | Eastern quoll | Dasyuromorphia | MgRas |
| This study | >DvMgRas5 | Eastern quoll | Dasyuromorphia | MgRas |
| This study | >DvMgRas6 | Eastern quoll | Dasyuromorphia | MgRas |
| This study | >DvMgRas7 | Eastern quoll | Dasyuromorphia | MgRas |
| This study | >DvMgRas8 | Eastern quoll | Dasyuromorphia | MgRas |
| This study | >DvMgRas9 | Eastern quoll | Dasyuromorphia | MgRas |
| This study | >Md_HRas | Grey short-tailed opossum | Didelphimorphia | Canonical |
| This study | >Md_KRas4B_DUP | Grey short-tailed opossum | Didelphimorphia | Canonical |

| Accession number | Gene | Species | Phylo_Group | Gene_Subfamily |
|---|---|---|---|---|
| This study | >Md_KRas4A | Grey short-tailed opossum | Didelphimorphia | Canonical |
| This study | >Md_KRas4B | Grey short-tailed opossum | Didelphimorphia | Canonical |
| This study | >Md_NRas | Grey short-tailed opossum | Didelphimorphia | Canonical |
| This study | >MdMgRas1 | Grey short-tailed opossum | Didelphimorphia | MgRas |
| This study | >MdMgRas2 | Grey short-tailed opossum | Didelphimorphia | MgRas |
| This study | >Me_HRas | Tammar Wallaby | Diprotodontia | Canonical |
| This study | >Me_KRas4A | Tammar Wallaby | Diprotodontia | Canonical |
| This study | >Me_KRas4B | Tammar Wallaby | Diprotodontia | Canonical |
| This study | >Me_NRas | Tammar Wallaby | Diprotodontia | Canonical |
| This study | >MeMgRas1 | Tammar Wallaby | Diprotodontia | MgRas |
| This study | >MeMgRas2 | Tammar Wallaby | Diprotodontia | MgRas |
| This study | >MeMgRas3 | Tammar Wallaby | Diprotodontia | MgRas |
| This study | >Ml_HRas | Greater bilby | Peramelemorphia | Canonical |
| This study | >Ml_KRas4a | Greater bilby | Peramelemorphia | Canonical |
| This study | >Ml_Kras4b | Greater bilby | Peramelemorphia | Canonical |
| This study | >Ml_NRas | Greater bilby | Peramelemorphia | Canonical |
| This study | >MlMgRas10 | Greater bilby | Peramelemorphia | MgRas |
| This study | >MlMgRas11 | Greater bilby | Peramelemorphia | MgRas |
| This study | >MlMgRas12 | Greater bilby | Peramelemorphia | MgRas |
| This study | >MlMgRas13 | Greater bilby | Peramelemorphia | MgRas |
| This study | >MlMgRas14 | Greater bilby | Peramelemorphia | MgRas |
| This study | >MlMgRas15 | Greater bilby | Peramelemorphia | MgRas |
| This study | >MlMgRas16 | Greater bilby | Peramelemorphia | MgRas |
| This study | >MlMgRas17 | Greater bilby | Peramelemorphia | MgRas |
| This study | >MlMgRas18 | Greater bilby | Peramelemorphia | MgRas |
| This study | >MlMgRas19 | Greater bilby | Peramelemorphia | MgRas |
| This study | >MlMgRas20 | Greater bilby | Peramelemorphia | MgRas |
| This study | >MlMgRas21 | Greater bilby | Peramelemorphia | MgRas |
| This study | >MlMgRas22 | Greater bilby | Peramelemorphia | MgRas |
| This study | >MlMgRas23 | Greater bilby | Peramelemorphia | MgRas |
| This study | >MlMgRas24 | Greater bilby | Peramelemorphia | MgRas |
| This study | >MlMgRas3 | Greater bilby | Peramelemorphia | MgRas |
| This study | >MlMgRas4 | Greater bilby | Peramelemorphia | MgRas |
| This study | >MlMgRas5 | Greater bilby | Peramelemorphia | MgRas |
| This study | >MlMgRas6 | Greater bilby | Peramelemorphia | MgRas |
| This study | >MlMgRas7 | Greater bilby | Peramelemorphia | MgRas |
| This study | >MlMgRas8 | Greater bilby | Peramelemorphia | MgRas |
| This study | >MlMgRas9 | Greater bilby | Peramelemorphia | MgRas |
| This study | >Pc_HRas | Koala | Diprotodontia | Canonical |
| This study | >Pc_KRas4A | Koala | Diprotodontia | Canonical |
| This study | >Pc_KRas4B | Koala | Diprotodontia | Canonical |
| This study | >Pc_NRas | Koala | Diprotodontia | Canonical |
| This study | >PcMgRas1 | Koala | Diprotodontia | MgRas |
| This study | >PcMgRas2 | Koala | Diprotodontia | MgRas |
| This study | >PcMgRas3 | Koala | Diprotodontia | MgRas |

| Accession number | Gene | Species | Phylo_Group | Gene_Subfamily |
|---|---|---|---|---|
| This study | >Pg_HRas | Eastern barred bandicoot | Peramelemorphia | Canonical |
| This study | >Pg_KRas4a | Eastern barred bandicoot | Peramelemorphia | Canonical |
| This study | >Pg_KRas4b | Eastern barred bandicoot | Peramelemorphia | Canonical |
| This study | >Pg_NRas | Eastern barred bandicoot | Peramelemorphia | Canonical |
| This study | >PgMgRas1 | Eastern barred bandicoot | Peramelemorphia | MgRas |
| This study | >PgMgRas10 | Eastern barred bandicoot | Peramelemorphia | MgRas |
| This study | >PgMgRas100 | Eastern barred bandicoot | Peramelemorphia | MgRas |
| This study | >PgMgRas101 | Eastern barred bandicoot | Peramelemorphia | MgRas |
| This study | >PgMgRas102 | Eastern barred bandicoot | Peramelemorphia | MgRas |
| This study | >PgMgRas103 | Eastern barred bandicoot | Peramelemorphia | MgRas |
| This study | >PgMgRas104 | Eastern barred bandicoot | Peramelemorphia | MgRas |
| This study | >PgMgRas105 | Eastern barred bandicoot | Peramelemorphia | MgRas |
| This study | >PgMgRas106 | Eastern barred bandicoot | Peramelemorphia | MgRas |
| This study | >PgMgRas107 | Eastern barred bandicoot | Peramelemorphia | MgRas |
| This study | >PgMgRas108 | Eastern barred bandicoot | Peramelemorphia | MgRas |
| This study | >PgMgRas109 | Eastern barred bandicoot | Peramelemorphia | MgRas |
| This study | >PgMgRas11 | Eastern barred bandicoot | Peramelemorphia | MgRas |
| This study | >PgMgRas110 | Eastern barred bandicoot | Peramelemorphia | MgRas |
| This study | >PgMgRas111 | Eastern barred bandicoot | Peramelemorphia | MgRas |
| This study | >PgMgRas112 | Eastern barred bandicoot | Peramelemorphia | MgRas |
| This study | >PgMgRas113 | Eastern barred bandicoot | Peramelemorphia | MgRas |
| This study | >PgMgRas114 | Eastern barred bandicoot | Peramelemorphia | MgRas |
| This study | >PgMgRas115 | Eastern barred bandicoot | Peramelemorphia | MgRas |
| This study | >PgMgRas116 | Eastern barred bandicoot | Peramelemorphia | MgRas |
| This study | >PgMgRas117 | Eastern barred bandicoot | Peramelemorphia | MgRas |
| This study | >PgMgRas12 | Eastern barred bandicoot | Peramelemorphia | MgRas |
| This study | >PgMgRas13 | Eastern barred bandicoot | Peramelemorphia | MgRas |
| This study | >PgMgRas14 | Eastern barred bandicoot | Peramelemorphia | MgRas |
| This study | >PgMgRas15 | Eastern barred bandicoot | Peramelemorphia | MgRas |
| This study | >PgMgRas16 | Eastern barred bandicoot | Peramelemorphia | MgRas |
| This study | >PgMgRas17 | Eastern barred bandicoot | Peramelemorphia | MgRas |
| This study | >PgMgRas18 | Eastern barred bandicoot | Peramelemorphia | MgRas |
| This study | >PgMgRas19 | Eastern barred bandicoot | Peramelemorphia | MgRas |
| This study | >PgMgRas2 | Eastern barred bandicoot | Peramelemorphia | MgRas |
| This study | >PgMgRas20 | Eastern barred bandicoot | Peramelemorphia | MgRas |
| This study | >PgMgRas21 | Eastern barred bandicoot | Peramelemorphia | MgRas |
| This study | >PgMgRas22 | Eastern barred bandicoot | Peramelemorphia | MgRas |
| This study | >PgMgRas23 | Eastern barred bandicoot | Peramelemorphia | MgRas |
| This study | >PgMgRas24 | Eastern barred bandicoot | Peramelemorphia | MgRas |
| This study | >PgMgRas25 | Eastern barred bandicoot | Peramelemorphia | MgRas |
| This study | >PgMgRas26 | Eastern barred bandicoot | Peramelemorphia | MgRas |
| This study | >PgMgRas27 | Eastern barred bandicoot | Peramelemorphia | MgRas |
| This study | >PgMgRas28 | Eastern barred bandicoot | Peramelemorphia | MgRas |
| This study | >PgMgRas29 | Eastern barred bandicoot | Peramelemorphia | MgRas |
| This study | >PgMgRas3 | Eastern barred bandicoot | Peramelemorphia | MgRas |
| This study | >PgMgRas30 | Eastern barred bandicoot | Peramelemorphia | MgRas |
| This study | >PgMgRas31 | Eastern barred bandicoot | Peramelemorphia | MgRas |
| This study | >PgMgRas32 | Eastern barred bandicoot | Peramelemorphia | MgRas |
| This study | >PgMgRas33 | Eastern barred bandicoot | Peramelemorphia | MgRas |

| Accession number | Gene | Species | Phylo_Group | Gene_Subfamily |
|---|---|---|---|---|
| This study | >PgMgRas34 | Eastern barred bandicoot | Peramelemorphia | MgRas |
| This study | >PgMgRas35 | Eastern barred bandicoot | Peramelemorphia | MgRas |
| This study | >PgMgRas36 | Eastern barred bandicoot | Peramelemorphia | MgRas |
| This study | >PgMgRas37 | Eastern barred bandicoot | Peramelemorphia | MgRas |
| This study | >PgMgRas38 | Eastern barred bandicoot | Peramelemorphia | MgRas |
| This study | >PgMgRas39 | Eastern barred bandicoot | Peramelemorphia | MgRas |
| This study | >PgMgRas4 | Eastern barred bandicoot | Peramelemorphia | MgRas |
| This study | >PgMgRas40 | Eastern barred bandicoot | Peramelemorphia | MgRas |
| This study | >PgMgRas41 | Eastern barred bandicoot | Peramelemorphia | MgRas |
| This study | >PgMgRas42 | Eastern barred bandicoot | Peramelemorphia | MgRas |
| This study | >PgMgRas43 | Eastern barred bandicoot | Peramelemorphia | MgRas |
| This study | >PgMgRas44 | Eastern barred bandicoot | Peramelemorphia | MgRas |
| This study | >PgMgRas45 | Eastern barred bandicoot | Peramelemorphia | MgRas |
| This study | >PgMgRas46 | Eastern barred bandicoot | Peramelemorphia | MgRas |
| This study | >PgMgRas47 | Eastern barred bandicoot | Peramelemorphia | MgRas |
| This study | >PgMgRas48 | Eastern barred bandicoot | Peramelemorphia | MgRas |
| This study | >PgMgRas49 | Eastern barred bandicoot | Peramelemorphia | MgRas |
| This study | >PgMgRas5 | Eastern barred bandicoot | Peramelemorphia | MgRas |
| This study | >PgMgRas50 | Eastern barred bandicoot | Peramelemorphia | MgRas |
| This study | >PgMgRas51 | Eastern barred bandicoot | Peramelemorphia | MgRas |
| This study | >PgMgRas52 | Eastern barred bandicoot | Peramelemorphia | MgRas |
| This study | >PgMgRas53 | Eastern barred bandicoot | Peramelemorphia | MgRas |
| This study | >PgMgRas54 | Eastern barred bandicoot | Peramelemorphia | MgRas |
| This study | >PgMgRas55 | Eastern barred bandicoot | Peramelemorphia | MgRas |
| This study | >PgMgRas56 | Eastern barred bandicoot | Peramelemorphia | MgRas |
| This study | >PgMgRas57 | Eastern barred bandicoot | Peramelemorphia | MgRas |
| This study | >PgMgRas58 | Eastern barred bandicoot | Peramelemorphia | MgRas |
| This study | >PgMgRas59 | Eastern barred bandicoot | Peramelemorphia | MgRas |
| This study | >PgMgRas6 | Eastern barred bandicoot | Peramelemorphia | MgRas |
| This study | >PgMgRas60 | Eastern barred bandicoot | Peramelemorphia | MgRas |
| This study | >PgMgRas61 | Eastern barred bandicoot | Peramelemorphia | MgRas |
| This study | >PgMgRas62 | Eastern barred bandicoot | Peramelemorphia | MgRas |
| This study | >PgMgRas63 | Eastern barred bandicoot | Peramelemorphia | MgRas |
| This study | >PgMgRas64 | Eastern barred bandicoot | Peramelemorphia | MgRas |
| This study | >PgMgRas65 | Eastern barred bandicoot | Peramelemorphia | MgRas |
| This study | >PgMgRas66 | Eastern barred bandicoot | Peramelemorphia | MgRas |
| This study | >PgMgRas67 | Eastern barred bandicoot | Peramelemorphia | MgRas |
| This study | >PgMgRas68 | Eastern barred bandicoot | Peramelemorphia | MgRas |
| This study | >PgMgRas69 | Eastern barred bandicoot | Peramelemorphia | MgRas |
| This study | >PgMgRas7 | Eastern barred bandicoot | Peramelemorphia | MgRas |
| This study | >PgMgRas70 | Eastern barred bandicoot | Peramelemorphia | MgRas |
| This study | >PgMgRas71 | Eastern barred bandicoot | Peramelemorphia | MgRas |
| This study | >PgMgRas72 | Eastern barred bandicoot | Peramelemorphia | MgRas |
| This study | >PgMgRas73 | Eastern barred bandicoot | Peramelemorphia | MgRas |
| This study | >PgMgRas74 | Eastern barred bandicoot | Peramelemorphia | MgRas |
| This study | >PgMgRas75 | Eastern barred bandicoot | Peramelemorphia | MgRas |
| This study | >PgMgRas76 | Eastern barred bandicoot | Peramelemorphia | MgRas |
| This study | >PgMgRas77 | Eastern barred bandicoot | Peramelemorphia | MgRas |
| This study | >PgMgRas78 | Eastern barred bandicoot | Peramelemorphia | MgRas |

| Accession number | Gene | Species | Phylo_Group | Gene_Subfamily |
|---|---|---|---|---|
| This study | >PgMgRas79 | Eastern barred bandicoot | Peramelemorphia | MgRas |
| This study | >PgMgRas8 | Eastern barred bandicoot | Peramelemorphia | MgRas |
| This study | >PgMgRas80 | Eastern barred bandicoot | Peramelemorphia | MgRas |
| This study | >PgMgRas81 | Eastern barred bandicoot | Peramelemorphia | MgRas |
| This study | >PgMgRas82 | Eastern barred bandicoot | Peramelemorphia | MgRas |
| This study | >PgMgRas83 | Eastern barred bandicoot | Peramelemorphia | MgRas |
| This study | >PgMgRas84 | Eastern barred bandicoot | Peramelemorphia | MgRas |
| This study | >PgMgRas85 | Eastern barred bandicoot | Peramelemorphia | MgRas |
| This study | >PgMgRas86 | Eastern barred bandicoot | Peramelemorphia | MgRas |
| This study | >PgMgRas87 | Eastern barred bandicoot | Peramelemorphia | MgRas |
| This study | >PgMgRas88 | Eastern barred bandicoot | Peramelemorphia | MgRas |
| This study | >PgMgRas89 | Eastern barred bandicoot | Peramelemorphia | MgRas |
| This study | >PgMgRas9 | Eastern barred bandicoot | Peramelemorphia | MgRas |
| This study | >PgMgRas90 | Eastern barred bandicoot | Peramelemorphia | MgRas |
| This study | >PgMgRas91 | Eastern barred bandicoot | Peramelemorphia | MgRas |
| This study | >PgMgRas92 | Eastern barred bandicoot | Peramelemorphia | MgRas |
| This study | >PgMgRas93 | Eastern barred bandicoot | Peramelemorphia | MgRas |
| This study | >PgMgRas94 | Eastern barred bandicoot | Peramelemorphia | MgRas |
| This study | >PgMgRas95 | Eastern barred bandicoot | Peramelemorphia | MgRas |
| This study | >PgMgRas96 | Eastern barred bandicoot | Peramelemorphia | MgRas |
| This study | >PgMgRas97 | Eastern barred bandicoot | Peramelemorphia | MgRas |
| This study | >PgMgRas98 | Eastern barred bandicoot | Peramelemorphia | MgRas |
| This study | >PgMgRas99 | Eastern barred bandicoot | Peramelemorphia | MgRas |
| This study | >Sc_Hras | Fat-tailed dunnart | Dasyuromorphia | Canonical |
| This study | >Sc_KRas4a | Fat-tailed dunnart | Dasyuromorphia | Canonical |
| This study | >Sc_KRas4b | Fat-tailed dunnart | Dasyuromorphia | Canonical |
| This study | >Sc_Nras | Fat-tailed dunnart | Dasyuromorphia | Canonical |
| This study | >ScMgRas1 | Fat-tailed dunnart | Dasyuromorphia | MgRas |
| This study | >ScMgRas10 | Fat-tailed dunnart | Dasyuromorphia | MgRas |
| This study | >ScMgRas11 | Fat-tailed dunnart | Dasyuromorphia | MgRas |
| This study | >ScMgRas12 | Fat-tailed dunnart | Dasyuromorphia | MgRas |
| This study | >ScMgRas13 | Fat-tailed dunnart | Dasyuromorphia | MgRas |
| This study | >ScMgRas14 | Fat-tailed dunnart | Dasyuromorphia | MgRas |
| This study | >ScMgRas2 | Fat-tailed dunnart | Dasyuromorphia | MgRas |
| This study | >ScMgRas3 | Fat-tailed dunnart | Dasyuromorphia | MgRas |
| This study | >ScMgRas4 | Fat-tailed dunnart | Dasyuromorphia | MgRas |
| This study | >ScMgRas5 | Fat-tailed dunnart | Dasyuromorphia | MgRas |
| This study | >ScMgRas6 | Fat-tailed dunnart | Dasyuromorphia | MgRas |
| This study | >ScMgRas7 | Fat-tailed dunnart | Dasyuromorphia | MgRas |
| This study | >ScMgRas8 | Fat-tailed dunnart | Dasyuromorphia | MgRas |
| This study | >ScMgRas9 | Fat-tailed dunnart | Dasyuromorphia | MgRas |
| This study | >Sh_DIRAS1 | Tasmanian devil | Dasyuromorphia | Ras subfamily |
| This study | >Sh_DIRAS2 | Tasmanian devil | Dasyuromorphia | Ras subfamily |
| This study | >Sh_GEM | Tasmanian devil | Dasyuromorphia | Ras subfamily |
| This study | >Sh_Hras | Tasmanian devil | Dasyuromorphia | Canonical |
| This study | >Sh_KRas4a | Tasmanian devil | Dasyuromorphia | Canonical |
| This study | >Sh_KRas4b | Tasmanian devil | Dasyuromorphia | Canonical |
| This study | >Sh_MRAS | Tasmanian devil | Dasyuromorphia | Ras subfamily |
| This study | >Sh_NKIRAS1 | Tasmanian devil | Dasyuromorphia | Ras subfamily |

| Accession number | Gene | Species | Phylo_Group | Gene_Subfamily |
|---|---|---|---|---|
| This study | >Sh_NKIRAS2 | Tasmanian devil | Dasyuromorphia | Ras subfamily |
| This study | >Sh_Nras | Tasmanian devil | Dasyuromorphia | Canonical |
| This study | >Sh_RAD | Tasmanian devil | Dasyuromorphia | Ras subfamily |
| This study | >Sh_RALA | Tasmanian devil | Dasyuromorphia | Ras subfamily |
| This study | >Sh_RALB | Tasmanian devil | Dasyuromorphia | Ras subfamily |
| This study | >Sh_RAP1A | Tasmanian devil | Dasyuromorphia | Ras subfamily |
| This study | >Sh_RAP1B | Tasmanian devil | Dasyuromorphia | Ras subfamily |
| This study | >Sh_RAP2A | Tasmanian devil | Dasyuromorphia | Ras subfamily |
| This study | >Sh_RAP2B | Tasmanian devil | Dasyuromorphia | Ras subfamily |
| This study | >Sh_RAP2C | Tasmanian devil | Dasyuromorphia | Ras subfamily |
| This study | >Sh_RASD1 | Tasmanian devil | Dasyuromorphia | Ras subfamily |
| This study | >Sh_RASL10A | Tasmanian devil | Dasyuromorphia | Ras subfamily |
| This study | >Sh_RASL10b | Tasmanian devil | Dasyuromorphia | Ras subfamily |
| This study | >Sh_RASL11A | Tasmanian devil | Dasyuromorphia | Ras subfamily |
| This study | >Sh_RASL11b | Tasmanian devil | Dasyuromorphia | Ras subfamily |
| This study | >Sh_RASL12 | Tasmanian devil | Dasyuromorphia | Ras subfamily |
| This study | >Sh_REM1 | Tasmanian devil | Dasyuromorphia | Ras subfamily |
| This study | >Sh_REM2 | Tasmanian devil | Dasyuromorphia | Ras subfamily |
| This study | >Sh_RERG | Tasmanian devil | Dasyuromorphia | Ras subfamily |
| This study | >Sh_RHEB | Tasmanian devil | Dasyuromorphia | Ras subfamily |
| This study | >Sh_RIT1 | Tasmanian devil | Dasyuromorphia | Ras subfamily |
| This study | >Sh_RIT2 | Tasmanian devil | Dasyuromorphia | Ras subfamily |
| This study | >Sh_RRAS | Tasmanian devil | Dasyuromorphia | Ras subfamily |
| This study | >Sh_RRAS2 | Tasmanian devil | Dasyuromorphia | Ras subfamily |
| This study | >ShMgRas1 | Tasmanian devil | Dasyuromorphia | MgRas |
| This study | >ShMgRas10 | Tasmanian devil | Dasyuromorphia | MgRas |
| This study | >ShMgRas11 | Tasmanian devil | Dasyuromorphia | MgRas |
| This study | >ShMgRas12 | Tasmanian devil | Dasyuromorphia | MgRas |
| This study | >ShMgRas13 | Tasmanian devil | Dasyuromorphia | MgRas |
| This study | >ShMgRas14 | Tasmanian devil | Dasyuromorphia | MgRas |
| This study | >ShMgRas16 | Tasmanian devil | Dasyuromorphia | MgRas |
| This study | >ShMgRas17 | Tasmanian devil | Dasyuromorphia | MgRas |
| This study | >ShMgRas2 | Tasmanian devil | Dasyuromorphia | MgRas |
| This study | >ShMgRas3 | Tasmanian devil | Dasyuromorphia | MgRas |
| This study | >ShMgRas4 | Tasmanian devil | Dasyuromorphia | MgRas |
| This study | >ShMgRas5 | Tasmanian devil | Dasyuromorphia | MgRas |
| This study | >ShMgRas6 | Tasmanian devil | Dasyuromorphia | MgRas |
| This study | >ShMgRas7 | Tasmanian devil | Dasyuromorphia | MgRas |
| This study | >ShMgRas8 | Tasmanian devil | Dasyuromorphia | MgRas |
| This study | >ShMgRas9 | Tasmanian devil | Dasyuromorphia | MgRas |

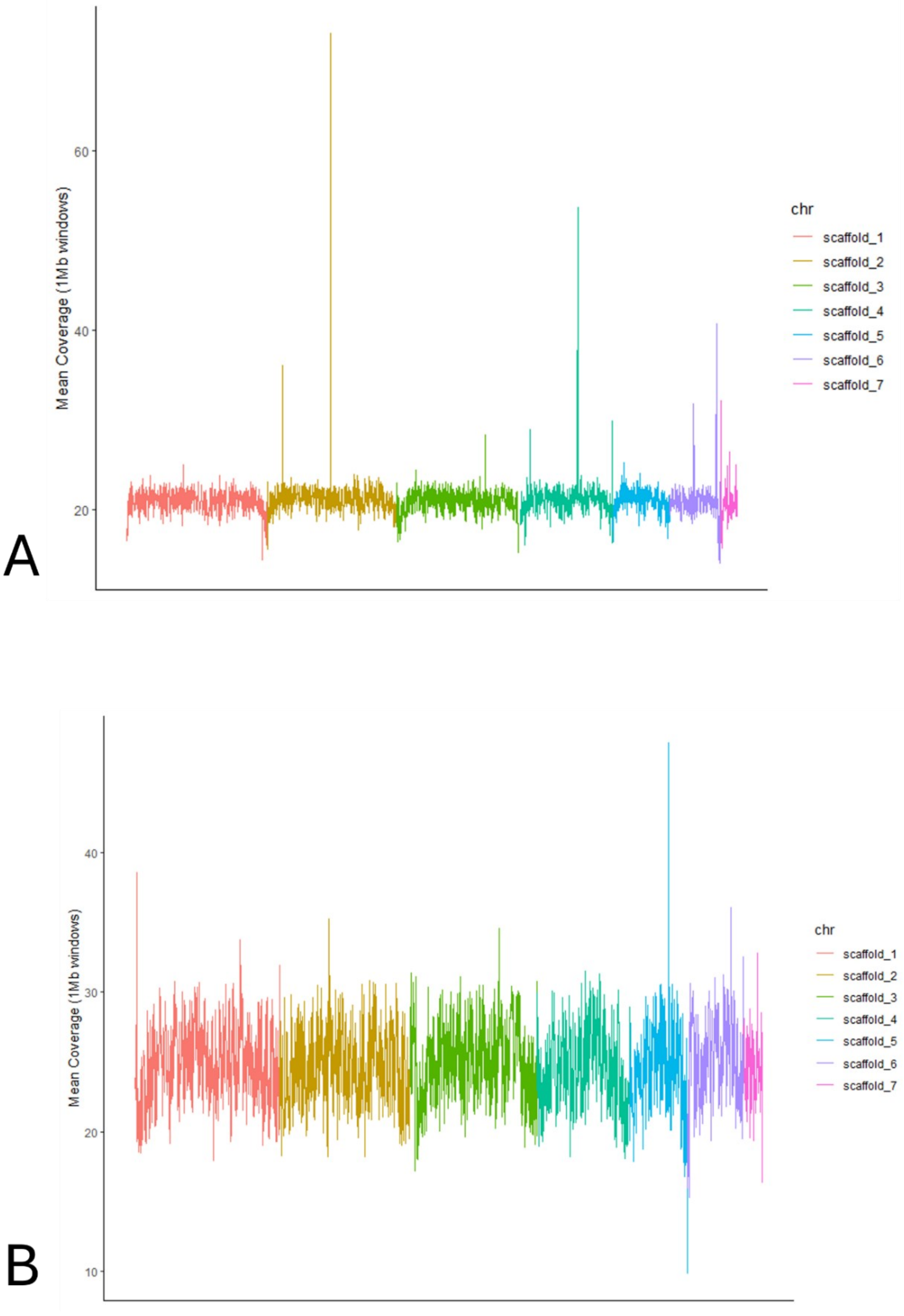


**Figure S1**. Plot showing mean coverage across the genome for A. Kowari and B. Bandicoot. Scaffolds are shown in different colours. For both species, coverage across the genome was approximately equal across all scaffolds, suggesting that both samples were from a homogametic individual.

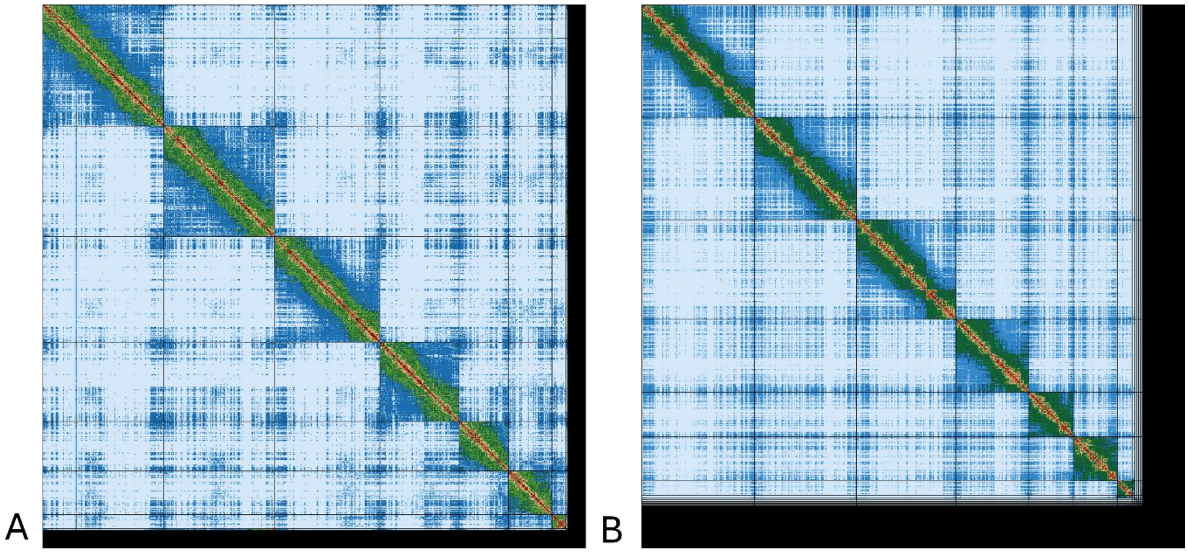

**Figure S2**. Hi-C contact map of the A. kowari and B. bandicoot assembly. Both indicate seven large scaffolds corresponding to the seven chromosomes expected in the species.

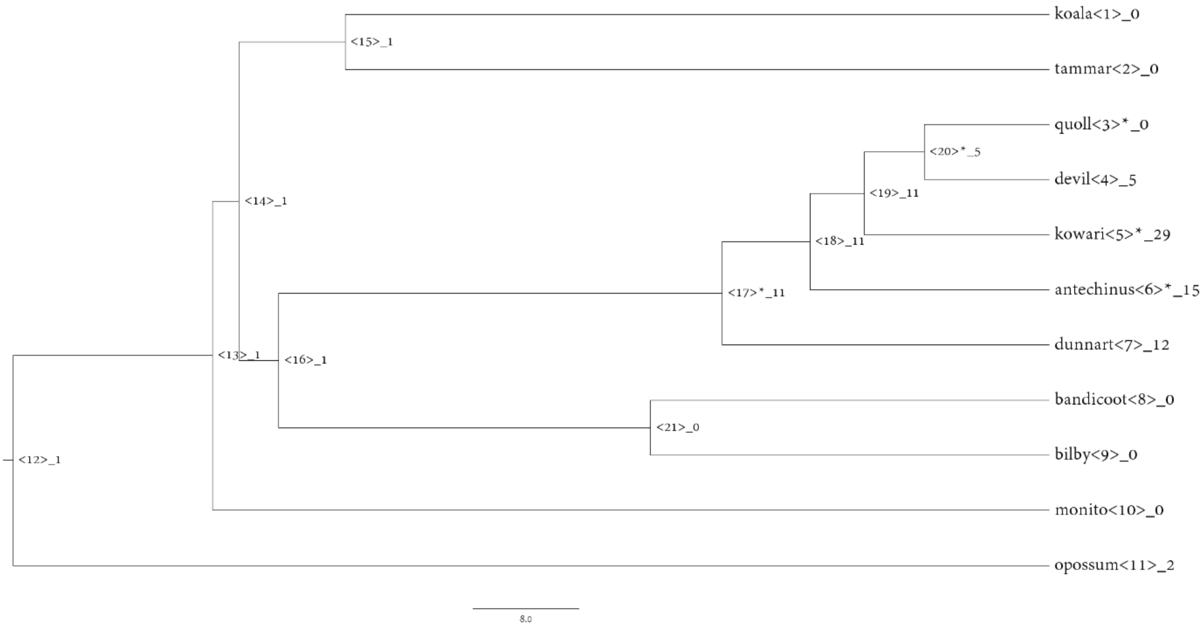


**Figure S3**. Gene tree for orthogroup HOG000767. The number after the underscore represents the number of genes each species has (or is predicted to have) in the orthogroup. Asterisk indicates that a statistically significant expansion or contraction occurred in this lineage. This orthogroup contains putative orthologs of *ATRX*, a tumour suppressor. The orthogroup underwent a significant expansion in the dasyurid ancestor but subsequent contractions in the ancestor of the quoll and devil.

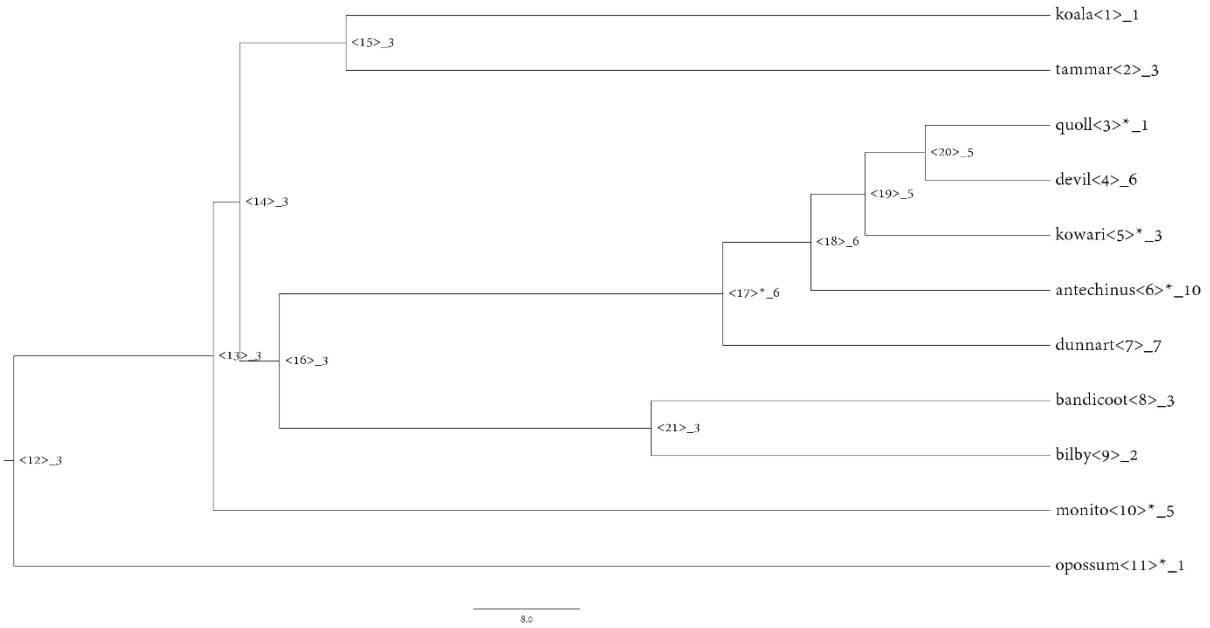


**Figure S4.** Gene tree for orthogroup HOG0001427. The number after the underscore represents the number of genes each species has (or is predicted to have) in the orthogroup. Asterisk indicates that a statistically significant expansion or contraction occurred in this lineage. This orthogroup contains copies of *SLC34A2*, a tumour supressor also involved in oncogenic fusion. The orthogroup underwent a significant expansion in the dasyurid ancestor but then subsequent contractions in the quoll and kowari.

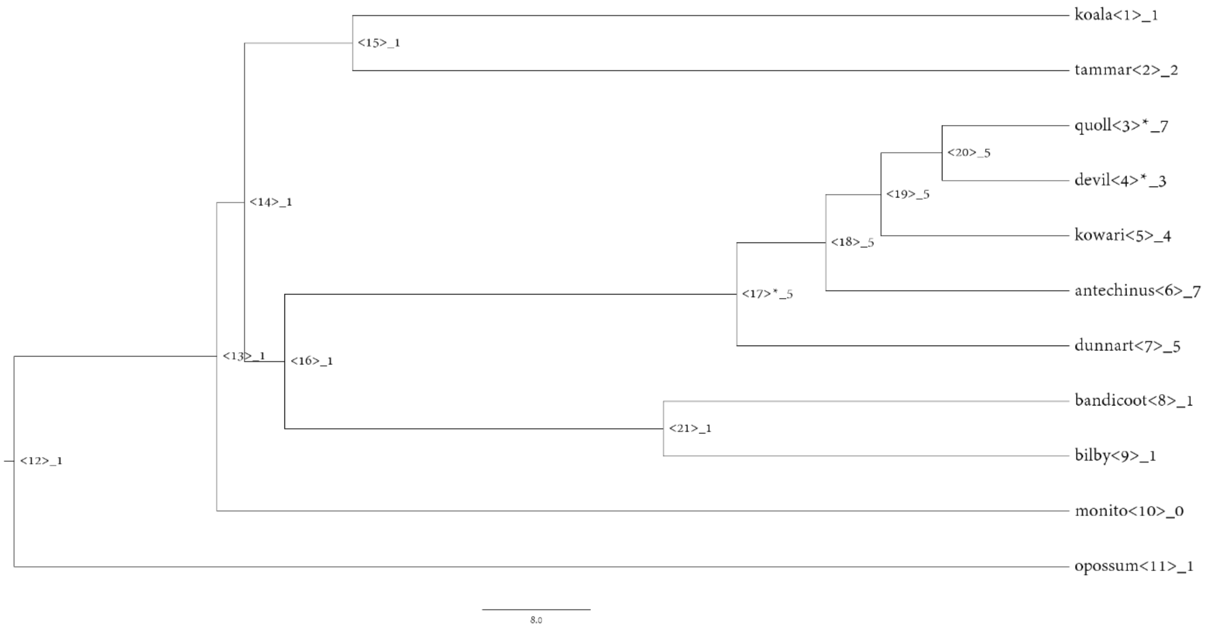


**Figure S5.** Gene tree for orthogroup HOG0002169. The number after the underscore represents the number of genes each species has (or is predicted to have) in the orthogroup. Asterisk indicates that a statistically significant expansion or contraction occurred in this lineage. This orthogroup contains copies of *TCEA1*, a gene involved in oncogenic fusions. The orthogroup underwent a significant expansion in the dasyurid ancestor but subsequently a contraction in the devil.

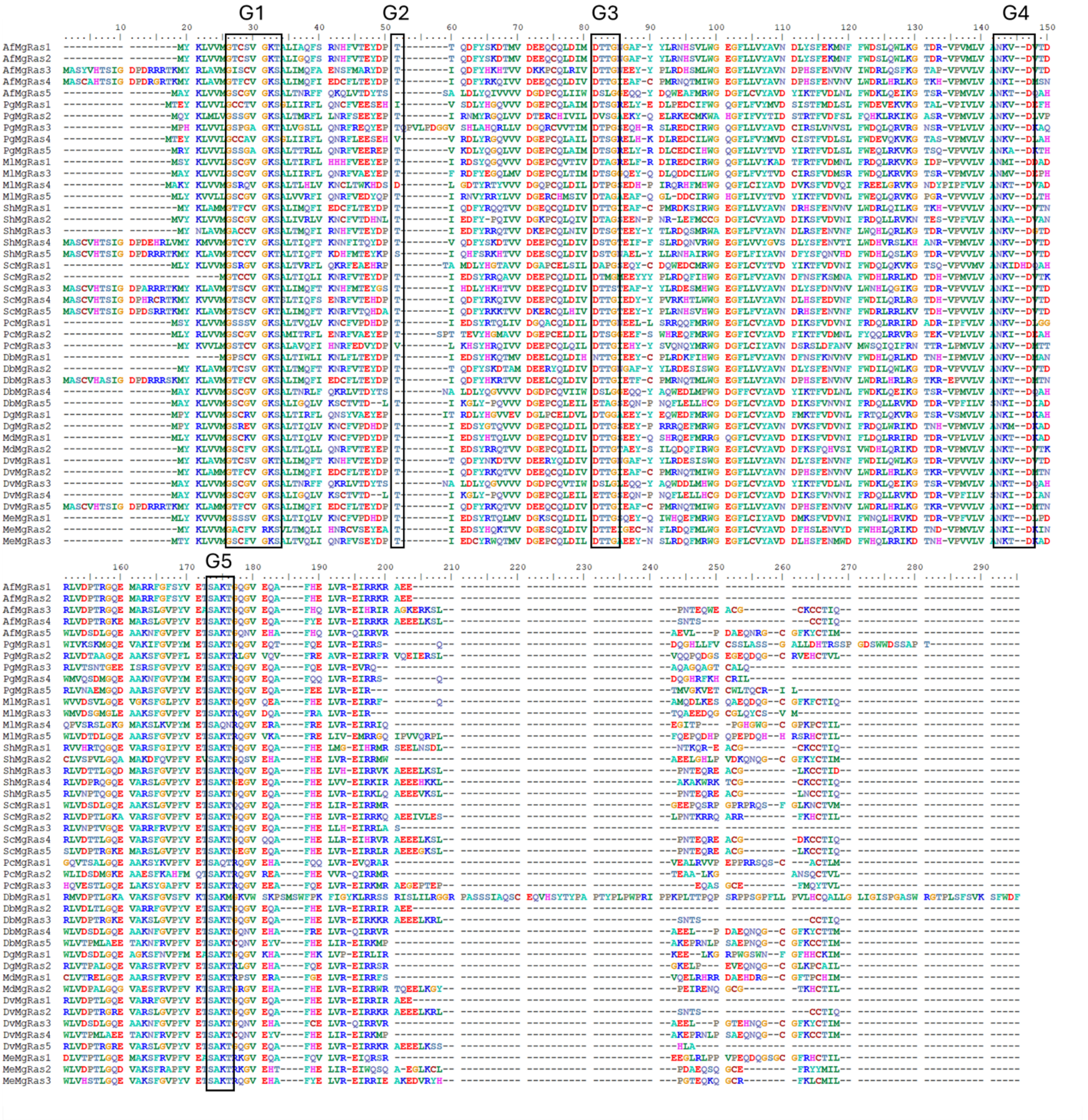


**Figure S6.** Multisequence alignment of MgRas genes with the five conserved G box motifs outlined in black. If a species had more than five genes, only the first five were represented in the figure. Species in the alignment are yellow-footed antechinus (Af), eastern barred bandicoot (Pg), bilby (Ml), Tasmanian devil (Sh), fat-tailed dunnart (Sc), koala (Pc), kowari (Db), monito del monte (Dg), grey short-tailed opossum (Md), eastern quoll (Dv) and tammar wallaby (Me).

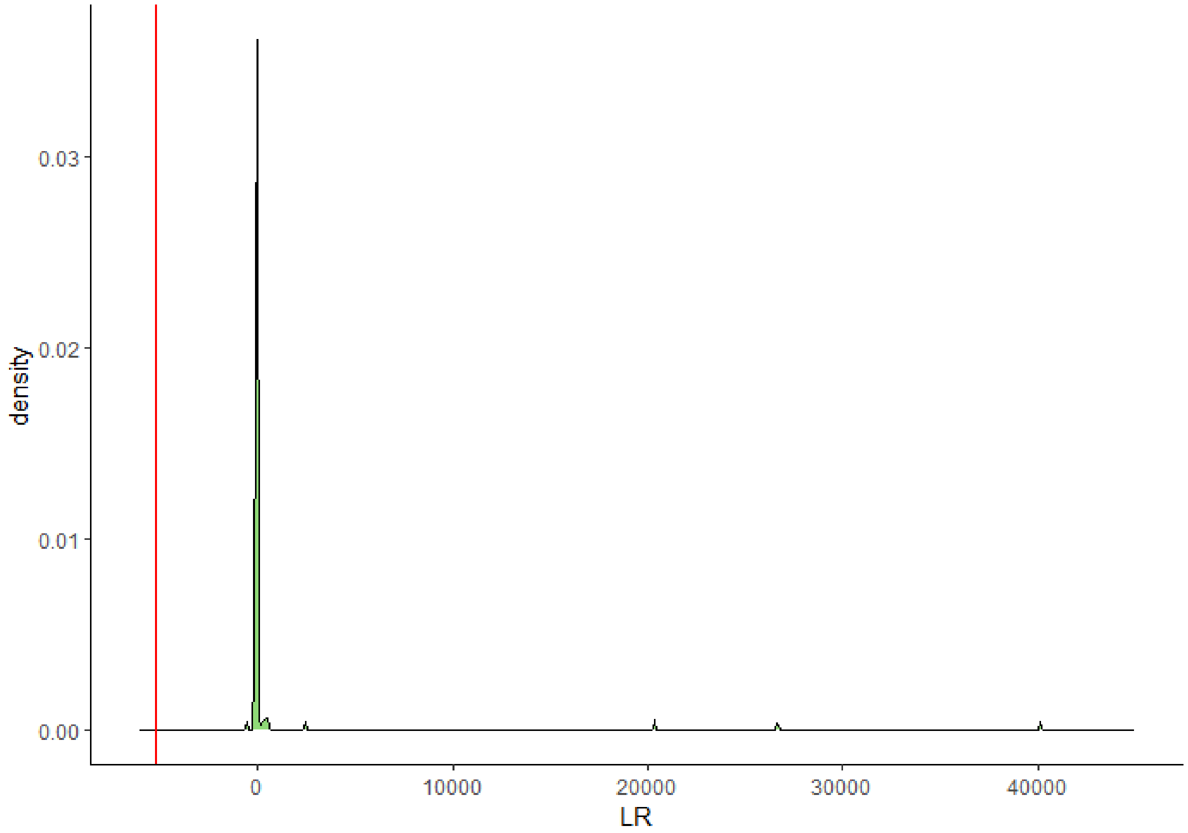


**Figure S7**. Likelihood ratio (LR) distribution under the null hypothesis. The green histogram represents the likelihood ratios obtained from the 100 simulations and the red line indicates the actual likelihood ratio. 0 of the 100 simulations had a value that was equal to or more extreme than the actual likelihood ratio, indicating that the probability (p-value) of obtaining this value under the null hypothesis is 0.

**References in the Supplementary Material**


Andrews, S. (2010). FastQC: a quality control tool for high throughput sequence data. In: Babraham Bioinformatics, Babraham Institute, Cambridge, United Kingdom.

Bolger, A. M., Lohse, M., & Usadel, B. (2014). Trimmomatic: a flexible trimmer for Illumina sequence data. *Bioinformatics*, *30*(15), 2114-2120.

Flynn, J. M., Hubley, R., Goubert, C., Rosen, J., Clark, A. G., Feschotte, C., & Smit, A. F. (2020). RepeatModeler2 for automated genomic discovery of transposable element families. *Proceedings of the National Academy of Sciences*, *117*(17), 9451-9457.

Haas, B. J., Papanicolaou, A., Yassour, M., Grabherr, M., Blood, P. D., Bowden, J., Couger, M. B., Eccles, D., Li, B., & Lieber, M. (2013). De novo transcript sequence reconstruction from RNA-seq using the Trinity platform for reference generation and analysis. *Nature protocols*, *8*(8), 1494-1512.

Kim, D., Paggi, J. M., Park, C., Bennett, C., & Salzberg, S. L. (2019). Graph-based genome alignment and genotyping with HISAT2 and HISAT-genotype. *Nature biotechnology*, *37*(8), 907-915.

Pertea, M., Pertea, G. M., Antonescu, C. M., Chang, T.-C., Mendell, J. T., & Salzberg, S. L. (2015). StringTie enables improved reconstruction of a transcriptome from RNA-seq reads. *Nature biotechnology*, *33*(3), 290-295.

Simão, F. A., Waterhouse, R. M., Ioannidis, P., Kriventseva, E. V., & Zdobnov, E. M. (2015). BUSCO: assessing genome assembly and annotation completeness with single-copy orthologs. *Bioinformatics*, *31*(19), 3210-3212.

Smit, A., Hubley, R., & Green, P. (2013-2015). *RepeatMasker Open-4.0*. In

Solovyev, V., Kosarev, P., Seledsov, I., & Vorobyev, D. (2006). Automatic annotation of eukaryotic genes, pseudogenes and promoters. *Genome biology*, *7*, 1-12.
